# Not all Notch pathway mutations are equal in the embryonic mouse retina

**DOI:** 10.1101/2023.01.11.523641

**Authors:** Bernadett Bosze, Julissa Suarez-Navarro, Illiana Cajias, Joseph A. Brzezinski, Nadean L Brown

## Abstract

In the vertebrate retina, combinations of Notch ligands, receptors, and ternary complex components determine the destiny of retinal progenitor cells by regulating *Hes* effector gene activity. Owing to reiterated Notch signaling in numerous tissues throughout development, there are multiple vertebrate paralogues for nearly every node in this pathway. These Notch signaling components can act redundantly or in a compensatory fashion during development. To dissect the complexity of this pathway during retinal development, we used seven germline or conditional mutant mice and two spatiotemporally distinct Cre drivers. We perturbed the Notch ternary complex and multiple *Hes* genes with two overt goals in mind. First, we wished to determine if Notch signaling is required in the optic stalk/nerve head for Hes1 sustained expression and activity. Second, we aimed to test if *Hes1, 3* and *5* genes are functionally redundant during early retinal histogenesis. With our allelic series, we found that disrupting Notch signaling consistently blocked mitotic growth and overproduced ganglion cells, but we also identified two significant branchpoints for this pathway. In the optic stalk/nerve head, sustained Hes1 is regulated independent of Notch signaling, whereas during photoreceptor genesis both Notch-dependent and -independent roles for *Rbpj* and *Hes1* impact photoreceptor genesis in opposing manners.

## INTRODUCTION

In vertebrate embryos, a central eye field is specified at the end of gastrulation and splits to form bilateral optic vesicles that evaginate from the ventral diencephalon. Multiple signaling pathways act to regionalize the growing optic vesicles, demarcating the optic stalk (OS), optic cup (OC) and retinal pigment epithelium (RPE) tissues. The OC gives rise to the neural retina, which is an excellent system for studying cell fate specification and differentiation. The retina is comprised of seven major cell classes that arise in a tightly controlled, but overlapping chronological order: retinal ganglion cells (RGCs), cone photoreceptors, horizontals, and a subset of amacrine neurons--before birth; and amacrines, rods, bipolars and Müller glia--mainly after birth. Throughout development, retinal progenitor cells (RPCs) balance their population size with generating neurons and glia [reviewed in 1, 2].

The deeply conserved Delta-Notch signaling pathway maintains the equilibrium between growth and differentiation in a myriad of tissues and often acts reiteratively within a single organ [3, 4]. In this transmembrane signaling system, ligand-receptor binding induces sequential proteolytic cleavages of the receptor protein to ultimately release the Notch intracellular domain (N-ICD), which forms a ternary complex with Rbpj (Recombination signaling binding protein, also termed CBF1) and Maml (Mastermind-like) (Fig 1A)[4]. These complexes bind DNA and transcriptionally activate target genes, including *Drosophila* ***H****airy* or ***E(s****pl)*, and the vertebrate *Hes* gene families [5-7]. Numerous studies have shown that too little or too much Notch signaling profoundly disrupts retinal neurogenesis (Suppl Table 1)[reviewed in 8]. The loss of Notch signaling from RPCs produces smaller retinas, with precocious retinal neurons [9-18]. Excess signaling blocks differentiation and induces RPC overproliferation [10], but at later stages promotes Müller glia, sometimes at the expense of differentiated neurons [19-22]. Notch signaling regulates multiple retinal cell processes, but is especially active during neurogenesis.

**Fig 1.**
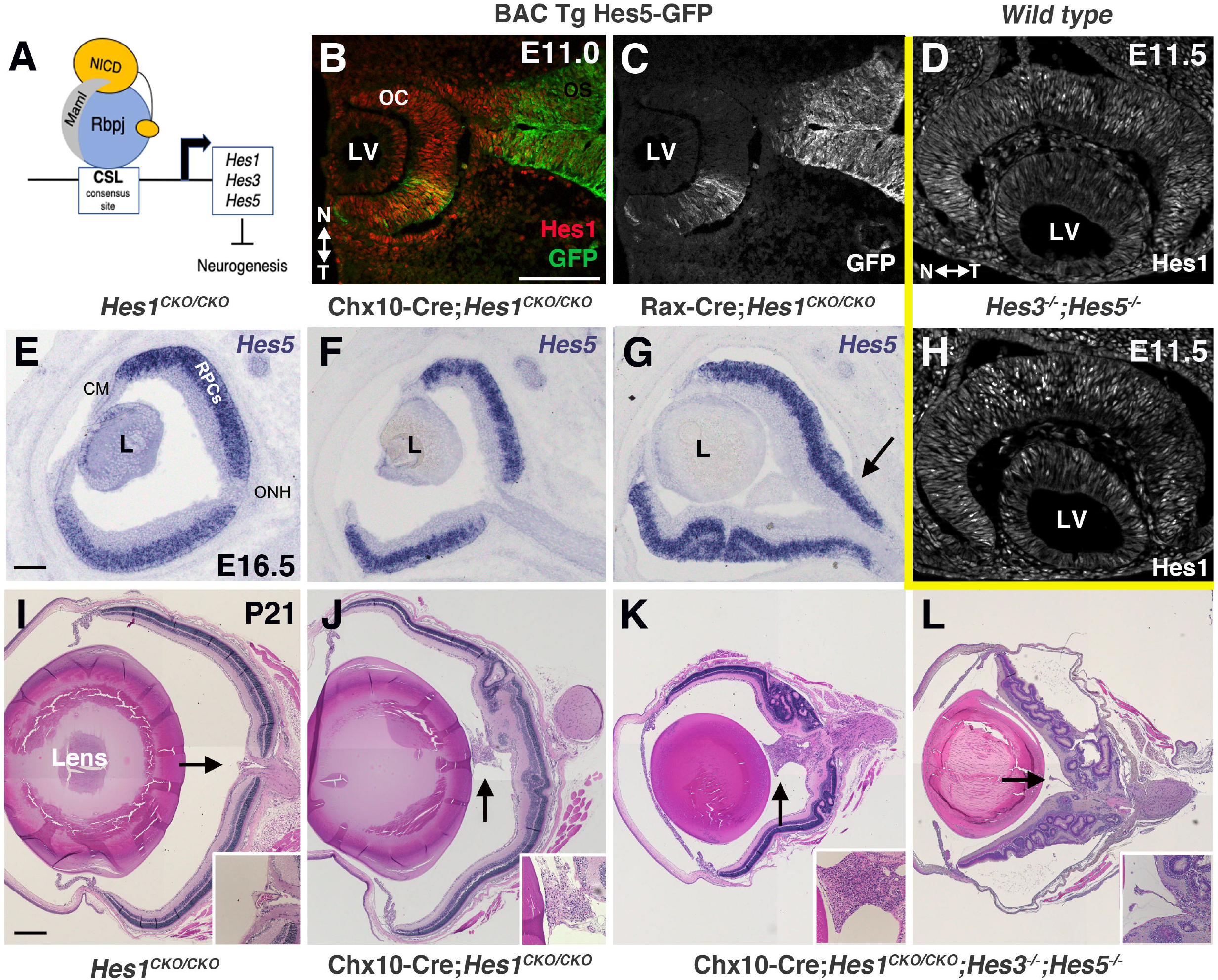
*Hes* gene relationships and mutant phenotypes. A) In the Notch pathway, ligand binding induces receptor protein cleavage that releases the intracellular domain (N-ICD), to form a ternary complex with Rbpj and Maml. These complexes bind DNA and transcriptionally activate *Hes* gene transcription. B-C) Hes1 + GFP immunolabeling show uniform Hes1 expression, but BAC Tg(*Hes5*-GFP) expression is restricted. D,H) At E11.5, Hes1 exhibits oscillating expression, which is unaffected in *Hes3*^*-/-*^*;Hes5*^*-/-*^ double mutants. E-G) *Hes5* mRNA inappropriately expands into the optic stalk (OS) when *Hes1* is conditionally removed with Rax-Cre (arrow in G), but not Chx10-Cre (F). I-L) A range of optic nerve head (ONH) defects in adult eyes (arrows all panels). When *Hes1* is absent there is an ectopic vitreal cell mass and sporadic retinal rosettes, yet Chx10-Cre;*Hes*^*TKO*^ eyes have extensive retinal lamination defects, abnormal ONH morphology (K) and ectopic vessels (arrow L). CSL= CBF1/Su(H)/Lag1; N= nasal; T = temporal; LV = lens vesicle; CM = ciliary margin; RPC = retinal progenitor cells. Bar in B = 10 microns, in E = 100 microns, in I = 200 microns; n ≥3 per age and genotype.

Most of the seven vertebrate *Hes* genes are Notch ternary complex targets (Fig 1a)[7, 23-26]. *Hes1, 3* and *5* are important in the nervous system, whereas *Hes2, 4* and *7* act in other parts of the body [7, 27]. The role of *Hes6* in development is debatable [reviewed in 7]. Both *Hes1* and *Hes5* can exhibit oscillating expression in neural progenitors or stem cells, controlling the balance between proliferation and differentiation [27]. For example, when *Hes1* is at high levels, progenitors remain proliferative, whereas low *Hes1* correlates with differentiation [28]. In the mouse spinal cord *Hes5* can be either sustained or oscillatory, with its frequency of oscillation correlating with onset of differentiation [29].

*Hes1* is an essential gene, whose loss causes prenatal lethality along with embryonic morphogenesis defects characterized by premature differentiation [30]. By comparison, complete loss of *Hes3* and/or *Hes5* has no impact on viability, but can induce discrete defects, suggesting that these paralogues are functionally redundant with *Hes1* in specific contexts. This is further supported by combinatorial *Hes* gene perturbation in other parts of the central nervous system (CNS), showing that *Hes1;Hes3;Hes5* triple mutants are the most severe [31-36].

The retinal-ONH/OS interface is one location where *Hes1* has a singular role, like at the brain isthmus [37, 38]. In both tissues, Hes1 protein is consistently high and sustained, unlike neural progenitor cells where it exhibits a variable oscillatory expression mode. It remains an open question whether Notch regulates sustained *Hes1* in the ONH/OS. Although *Hes* transcriptional repressors are frequent Notch downstream effectors, their activities can also be guided by other pathways, such as Shh signaling [39]. During retinal neurogenesis, loss of *Hes1* results in the overproduction of RGCs [15, 30, 38, 40], but also a decrease in cone photoreceptors (Supplementary Table 1) [38]. This does not align with other Notch signaling pathway mutants (*e*.*g*., *Dll1/4, Notch1*, or *Rbpj*), whose loss each overproduces both RGCs and cones (Supplementary Table 1). This misalignment suggests that although *Hes1* remains under Notch regulation during retinal neurogenesis, it is also regulated independently by other signaling pathways.

To dissect how *Hes1* functions in the retina independently of Notch signaling, we examined the prenatal retinal phenotypes of single and combined *Hes* mutant mice [31, 41]. Because *Hes* triple germline mutants die soon after gastrulation, a *Hes1* conditional mutation (*Hes1*^*CKO/CKO*^) was combined with *Hes3*^*-/-*^*;Hes5*^*-/-*^ germline mutant alleles, to effectively generate tissue-specific *Hes* triple mutants (*Hes*^*TKO*^) [31, 41]. To understand the extent to which *Hes*^*TKO*^ phenocopies other Notch pathway phenotypes, we simultaneously evaluated two essential ternary complex components, by generating *Rbpj*^*CKO/CKO*^ and *ROSA*^*dnMaml-GFP/+*^ retinal mutants. The *ROSA*^*dnMaml-GFP/+*^ allele is under flox-stop control, and dominantly creates inactive Notch transcriptional complexes, using a truncated Mastermind-nGFP fusion protein that binds to endogenous N-ICD and *Rbpj* [42-45]. We used two Cre drivers (Rax-Cre and Chx10-Cre), for all loss-of-function alleles generated, taking advantage of spatially overlapping, but temporally offset Cre activation, to tease apart morphologic versus neurogenic roles for these genes [38, 46]. This also allowed for direct comparison with previous findings in the mouse retina (Supplemental Table 1) [9-18].

We determined that sustained Hes1 expression in ONH/OS cells is Notch-independent, but within the adjacent retinal compartment, *Hes1* and *Hes5* are functionally redundant during the initiation of neurogenesis. We directly compared *Hes*^*TKO*^ versus *Rbpj* conditional mutants and found that *Hes* functions are consistent with Notch signaling during RPC growth, onset of RGC neurogenesis and apoptotic cell death. We also discovered that Maml cofactor activities are not exclusive to the Notch ternary complex, since *ROSA*^*dnMaml-GFP/+*^ retinal mutants displayed unique nasal-temporal patterning defects. Our phenotypic analyses uncovered that both Notch-dependent and -independent functions impact the onset of photoreceptor genesis, particularly highlighting the opposing roles of *Rbpj* and *Hes1* during cone photoreceptor development. Although *Hes*^*TKO*^ mutants rescued the loss of cones found in *Hes1* single mutants, they did not phenocopy excess cone formation seen in *Notch1* or *Rbpj* mutants [11, 15, 17, 18]. We conclude that an additional, unknown level of genetic redundancy (likely to be Notch-independent) occurs at the onset of cone photoreceptor neurogenesis. Our findings highlight the initiation of photoreceptor development as an important convergence point for integrating the inputs from multiple signaling pathways.

## RESULTS

During mouse nervous system development, *Hes1* is present in the anterior neural plate, optic vesicle and optic cup, several days prior to the onset of retinal neurogenesis [30, 47]. At these early stages, *Hes1* mRNA and protein are uniformly expressed. As the first cohort of optic cup cells exit mitosis and differentiate into neurons, mitotic RPCs switch to an oscillating mode of *Hes1* expression. However, optic nerve head (ONH) and optic stalk (OS) cells sustain uniform Hes1 expression [38]. *Hes5* mRNA appears after *Hes1* in the optic cup RPCs, prior to the formation of RGC neurons. *Hes5* is also found in the diencephalon, but not in E10.5-E13 optic stalk cells (data not shown). Although *Hes3* is active in the CNS as early as E9.5 [48], we could not detect it in the retina prior to E18 (not shown). To understand the functional redundancy of *Hes* genes during retina-ONH boundary formation and early retinal histogenesis, we directly compared the expression of and loss of multiple *Hes* genes during embryonic eye development.

The mouse Hes5-GFP BAC transgene is an accurate reporter of *Hes5* expression, enabling direct correlation of Hes1 and Hes5-GFP expression during development [49]. At E10-E11, Hes5-GFP is patchy in the optic cup with its biggest domain on the temporal side (Fig 1B,C). Hes5-GFP is also present in the diencephalon (Fig 1B,C), but does not overlap with Pax2 in the forming visual system (data not shown). This contrasts Hes1, present in all optic cup and stalk cells (Fig 1B,C). When the Hes5-GFP BAC transgene was previously placed on a *Hes1*^*-/-*^ background, GFP expression was derepressed throughout the optic cup and stalk, consistent with *Hes1* suppression of *Hes5* in other neural tissues [34, 49, 50]. *Hes3* functionally overlaps with *Hes1* in the brain isthmus [48], so we also evaluated the *Hes3* germline mutation, to understand if potentially low levels of *Hes3* may functionally overlap with *Hes1* in the ONH. Next, we tested whether Hes1 is depends on other *Hes* genes. *Hes3* and *Hes5* are <1Mb apart on mouse chromosome 4, and their knockout alleles are transmitted together as one mutant haplotype (Suppl Table 2) [31, 37]. We examined Hes1 ocular expression from E10.5-E16.5 within *Hes3*^*-/-*^*;Hes5*^*-/-*^ mice, and found the early uniform, oscillating RPC, and sustained ONH/OS Hes1 domains are all normal (Figs 1D,1H, 2C and data not shown). Thus, *Hes1* is not cross-regulated by either *Hes3* or *Hes5*. We also confirmed that *Hes3*^*-/-*^*;Hes5*^*-/-*^ mutants have normal retinal morphology and cell-type composition across eight developmental stages (E10.5-P21) (Suppl Fig 1 and data not shown). Next, we checked for reciprocal regulation by evaluating *Hes5* mRNA in E13.5 or E16.5 *Hes1* conditional (CKO) mutants, using two Cre drivers whose activation is temporally offset (Rax-Cre versus Chx10-Cre) [38, 46]. *Hes5* mRNA was unaffected in Chx10-Cre;*Hes1*^*CKO/CKO*^ retinas (Fig 1F), whereas with earlier deletion using Rax-Cre, *Hes5* mRNA abnormally extended into the optic stalk (arrow in Fig 1G). We conclude *Hes1* normally suppresses *Hes5* as the ONH boundary forms between E12-E13. While this outcome is consistent with direct cross-regulation, the expansion of the *Hes5* domain could also be due to retinal tissue overgrowth and displacement of the ONH/OS boundary.

To understand if the loss of multiple *Hes* genes is more catastrophic than *Hes1* alone, we used the same two Cre drivers with *Hes*^*TKO*^ mice (*Hes1*^*CKO/CKO*^; *Hes3*^*-/-*^*;Hes5*^*-/-*)^. We collected litters at E11, E13.5, E16.5, P0 (birth) and P21 (Suppl Table 2). Rax-Cre;*Hes*^*TKO*^ mutants were not viable beyond E13 and displayed more severe phenotypes than *Hes1* single mutants (Suppl Table 2, Figs 3, 4)[38]. Next, we directly compared P21 Chx10-Cre;*Hes*^*TKO*^ to Chx10-Cre;*Hes1*^*CKO/CKO*^ mutant eyes (n=3 biologic replicates/genotype). *Hes1* single mutants have defective retinal lamination, rosettes, and occasionally a small, vitreal cell mass (Fig 1J arrow). By contrast, adult Chx10-Cre;*Hes*^*TKO*^ eyes have more severe microphthalmia, retinal lamination and rosetting defects, and fully-penetrant vascularized tissue in the vitreous (Figs 1K,1L). In some sections, ectopic blood vessels protruded from the ONH (Figs 1K,1L arrows). We conclude that there are specific contexts in the developing retina where *Hes* genes are functionally redundant. We therefore initiated a deeper phenotypic evaluation at ages when both triple mutants are viable.

### *Hes*^*TKO*^ and *Rbpj* mutants are more severe than *dnMaml*

In theory, the functional disruption or loss of *Rbpj* or *Maml* from the Notch ternary complex should be as severe as removing major transcriptional targets like the *Hes* genes. At E13.5 *Hes5* mRNA is normally restricted to RPCs (Fig 2B). We validated the loss of *Hes5* mRNA in each allelic combination containing *Hes5*^*-/-*^ homozygotes (Figs 2D, 2F, 2H; n=3 mutants). We also verified that *Hes3* mRNA is completely missing from its early CNS domains in allelic combinations that include *Hes3*^*-/-*^ (not shown). Then, we analyzed both variable and sustained Hes1 patterns after Cre-mediated removal of all three *Hes* genes (Fig 2). In E13.5 Rax-Cre;*Hes*^*TKO*^ eyes, RPC and ONH/OS cells are devoid of Hes1 protein (Fig 2E). Because Chx10-Cre activates later in a retinal-restricted lineage [38], we expected Hes1 removal in RPC, but not ONH/OS cells. However, in E13.5 Chx10-Cre;*Hes*^*TKO*^ eyes, Hes1 clearly persists in both domains (Fig 2G). We hypothesized this is due to mosaic Chx10-Cre expression [46, 51]. To explore this idea further, we directly compared Rax-Cre versus Chx10-Cre deletion of *Rbpj* (Suppl. Fig 2; n=3 replicates for age and genotype). As expected, E13.5 Rax-Cre;*Rbpj*^*CKO/CKO*^ mutants had a cell autonomous loss of *Rbpj* from RPC, ONH/OS, and RPE cells (compare Suppl Figs 2A, 2B,2B’). Although Hes1 was absent from the optic cup and RPE (compare Suppl Figs 2A’, 2B”), ONH/OS cells still express it, demonstrating that sustained Hes1 is independent of Notch signaling.

**Fig 2.**
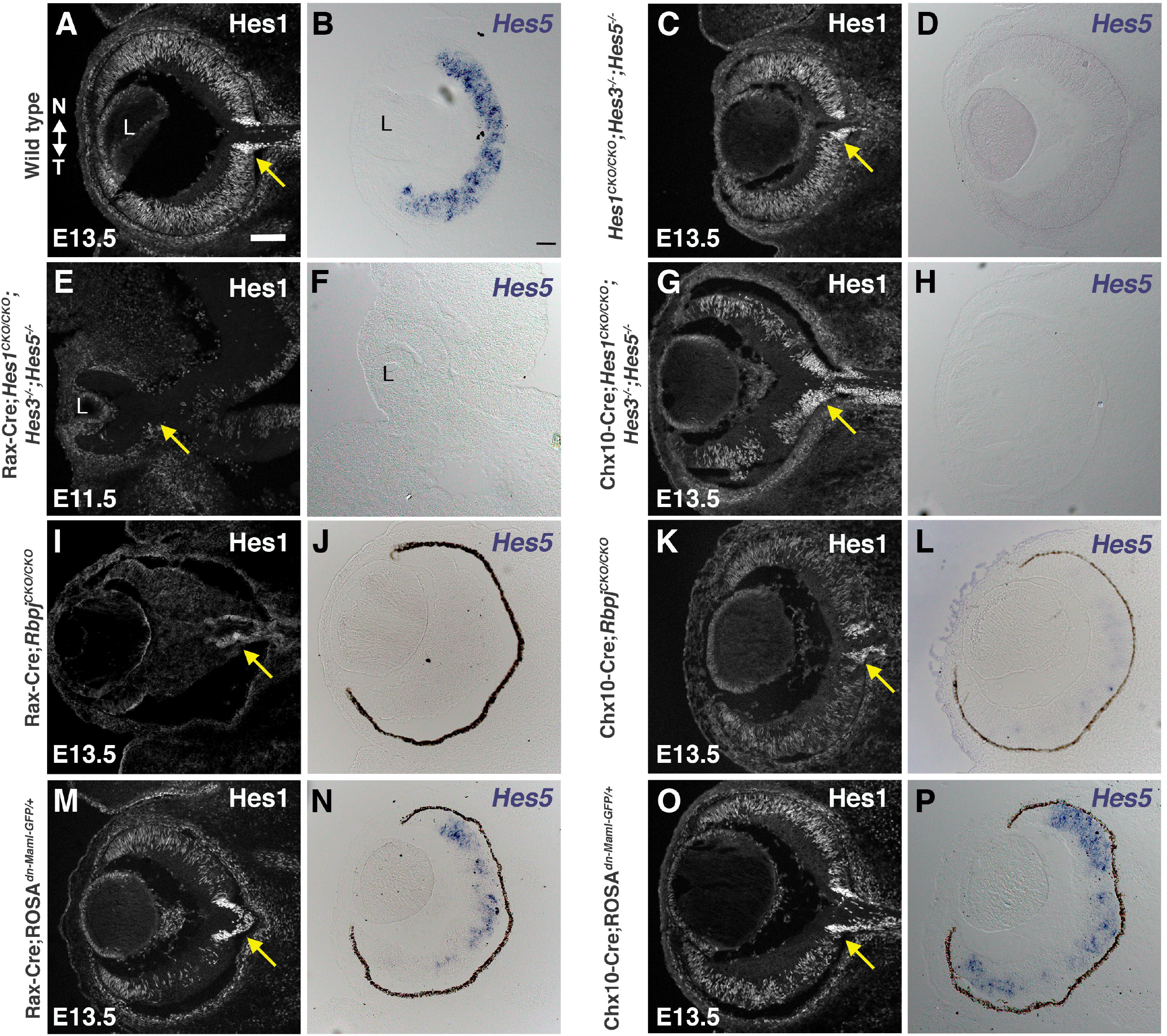
*Hes1* and *Hes5* expression in *Rbpj*, dnMaml and *Hes* triple retinal mutants. (A,C,E,G,I,K,M,O) Anti-Hes1 labeling of E11.5 or E13.5 cryosections. Hes1 is missing in Rax-Cre;*Hes*^*TKO*^ and Rax-Cre;*Rbpj*^*CKO/CKO*^ RPCs (E, I), with the intense Hes1+ ONH domain (yellow arrows) only lost in Rax-Cre;*Hes*^*TKO*^ eyes. (B,D,F,H,J,L,N,P) *Hes5* mRNA is missing in all *Hes5* germline mutants. (F,H) Both *Rbpj* conditional mutants effectively block *Hes5* mRNA expression (J,L). dnMaml partially knocks down Hes1 (M,O) and *Hes5* (N,P), with temporal optic cup more affected in Rax-Cre;*ROSA*^*dnMaml1-GFP/+*^ eyes. All panels oriented nasal up (noted in A); L = lens; scalebar in A = 100 microns, B = 50 microns, n= 3/3 mutants per genotype.

We took advantage of a Cre-GFP fusion protein within the Chx10-Cre driver to directly compared GFP and Rbpj coexpression in Chx10-Cre;*Rbpj*^*CKO/CKO*^ and control Chx10-Cre;*Rbpj*^*CKO/+*^ retinal sections (Suppl Fig 2C, 2E, 2G, 2I; n=3 biologic replicates/genotype). This Chx10-Cre BAC transgene encodes a Cre-GFP fusion protein, allowing for test of cell autonomy in the GFP cell population [46]. At E13.5 we noted a strong knockdown of Rbpj protein (Suppl Figs 2C” vs 2E”), yet at E16.5 there were proportionally more Rbpj-expressing retinal cells that also lacked GFP, identifying them as wild type (compare Suppl Fig 2G’ to 2I’). Hes1 was not obviously downregulated at either age (Suppl Figs 2D,2F,2H,2J). We concluded that Chx10-Cre phenotypes generated through E13.5 are informative, but beyond this stage the wild type cohort (GFP-neg) outcompetes mutant (GFP+) cells [52, 53], providing ample levels of Notch signaling. Evaluation of *Hes5* mRNA further confirmed Rax-Cre is the more effective driver, since we detected *Hes5* in Chx10-Cre;*Rbpj*^*CKO/CKO*^ retinas (compare Figs 2J to 2L). So, we confined subsequent phenotypic analyses to E13.5, when Rax-Cre mutants are viable and Chx10-Cre mosaicism is less impactful. Next, we examined *Hes1* and *Hes5* expression in E13.5 Rax-Cre;*ROSA*^*dnMAML-GFP/+*^ and Chx10-Cre;*ROSA*^*dnMAML-GFP/+*^ retinas. Both *Hes* genes are still clearly expressed, although there was a stronger knockdown in the Rax-Cre;*ROSA*^*dnMAML-GFP/+*^ temporal retina (Figs 2M-2P). We presume this *dnMAML* allele exhibits only a partial dominant negative effect in the developing eye, but analyzed it further to learn when, where and the degree to which it mimics *Rbpj*^*CKO/CKO*^ and *Hes*^*TKO*^ mutants.

### Early loss of Notch signaling does not impact optic cup patterning

The optic vesicle and cup are patterned along dorsal-ventral (D/V) and nasal-temporal (N/T) axes. *Hes1* mutants were already known to have no D/V ocular phenotypes [38, 47]. We checked for mispatterning of the N/T axis, since Pax2 is displaced in Rax-Cre; *Hes1*^*CKO/CKO*^ eyes, and *Pax2* mutants eyes contain such defects [54]. We compared the nasal-restricted marker Foxg1 [55, 56] among the six Rax-Cre or Chx10-Cre-induced mutants at E13.5 and E16.5 (Suppl Fig 3; n=3 replicates/ age + genotype). We saw normal retinal expression, with two exceptions. In E13.5 Rax-Cre;*Hes*^*TKO*^ eyes (Suppl Fig 3D) Foxg1 had spread into the nasal optic stalk, and in E16.5 Rax-Cre;*ROSA*^*dnMAML-GFP/+*^ mutants it was mislocalized to the temporal retina and subretinal space (arrow in Suppl Fig 3J), a cell-free zone between the apical retina and RPE. We presume the displaced cells are RPCs that appear in the subretinal space in some Notch pathway mutants due to loss of the outer limiting membrane from the apical surface of the optic cup [57, 58].

The optic cup is split into retina, rpe, stalk, ciliary tissue after DV/NT patterning of the retina. For example, the future neural retina becomes surrounded by a monolayer of nonneuronal cells, the retinal pigmented epithelium (RPE). Vsx2/Chx10 (RPCs) and Mitf (RPE) transcription factors delineate these tissues, and then actively maintain this boundary [59-61]. We compared Vsx2 and Mitf expression among all six E13.5 mutants (Figs 3A-3G), expecting there would be fewer RPCs. All E13.5 Rax-Cre-generated mutants had noticeably smaller eyes (Figs 3B,3C,3D), but Chx10-Cre generated mutants were typically of normal size (Fig 3E-G). For all six allelic combinations, the RPE formed correctly, but in Rax-Cre;*Hes*^*TKO*^ eyes this tissue extended into the optic stalk (Fig 3D), phenocopying Rax-Cre; *Hes1*^*CKO/CKO*^ mutants [38]. Rax-Cre;*Rbpj*^*CKO/CKO*^ and Rax-Cre;*Hes*^*TKO*^ mutants displayed the same RPC defect, namely patches of Vsx2-negative cells in the proximal optic cup, where neurogenesis normally initiates (Figs 3B,3D). We conclude that Notch signaling has no overt role in D/V and N/T patterning, or retinal/RPE specification (Fig 3B).

**Fig 3.**
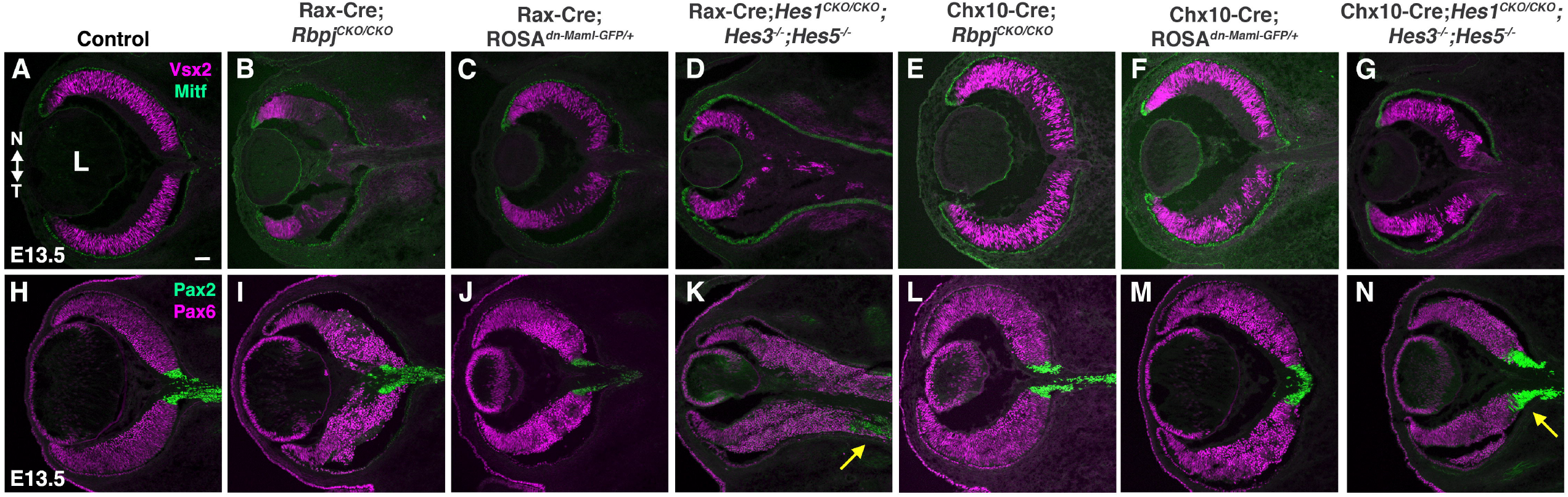
Ocular tissue patterning defects in Notch pathway mutants. (A-G) Vsx2 + Mitf immunolabeling marks the retinal-RPE boundary at E13.5. Vsx2+ RPCs were disorganized in the all mutants, but there is a smaller domain only in Rax-Cre;*Rbpj*^*CKO/CKO*^ and Rax-Cre;*Hes*^*TKO*^ eyes (B,D). (H-N) Pax6-Pax2 colabeling delineates the retinal-optic stalk boundary. Rax-Cre;*Hes*^*TKO*^ eyes (arrow in K) had the greatest retinal tissue expansion at the expense of Pax2+ ONH/optic stalk. The Pax2 domain was misshapen only in Chx10-Cre;*Hes*^*TKO*^ eyes (arrow in N). L = lens; all panels are oriented nasal up (noted in A); n ≥3 biologic replicates/genotype scalebar = 50 microns.

At E12, another tissue boundary forms between the neural retina and optic stalk, establishing a ring of cells called the optic nerve head (ONH). This boundary is delineated by the abutting expression of the transcription factors Pax6 (RPCs) and Pax2 (ONH/OS)[62]. Although the molecular mechanisms regulating this boundary are not well understood, its formation requires *Hes1* and *Pax2* activities [38, 54, 62]. To understand whether Notch signaling has a role here, we performed Pax6/Pax2 colabeling at E13.5 among all mutants (Figs 3I-3N). The Rax-Cre;*Hes*^*TKO*^ eyes, although not phenotypically different than Rax-Cre;*Hes1*^*CKO/CKO*^ mutants, were the most severe, with Pax6+ retinal tissue in the optic stalk territory, displacing the Pax2 domain (arrow in Fig 3K). Although the Pax6-Pax2 boundary is intact in Rax-Cre;*Rbpj*^*CKO/CKO*^ eyes, ONH shape was attenuated compared to controls (Fig 3I). Interestingly, the proximal-most optic cup cells, those lacking Vsx2, still express Pax6 (compare Figs 3B to 3I), suggesting these cells had differentiated into neurons (see Fig 5). The Rax-Cre;*ROSA*^*dnMAML-GFP/+*^ eyes were largely unaffected, although ONH shape was abnormal (Fig 3J). In all three Chx10-Cre generated mutants, a Pax6-Pax2 boundary is clearly discernable (Figs 3L-3N). But for Chx10-Cre;*Hes*^*TKO*^ mutants, there was a unique presence of ectopic Pax2 within the retinal territory (arrow in Fig 3N), demonstrating overlapping *Hes* gene function at this boundary (Figs 3D,3K,3N).

The ONH and brain isthmus share multiple features, including Pax2 and sustained Hes1 expression [37, 48, 63]. Isthmus cells proliferate at a slower rate, but this feature not been explored for the ONH. So, we examined Cyclin D2 (Ccnd2) expression, which is expressed by glial brain cells and intermediate neural progenitors with slow cycling kinetics [64, 65]. Interestingly, E13.5 ONH cells normally express Ccnd2, which is also downstream of Notch signaling in the ocular lens [66, 67]. Ccnd2 was downregulated in Rax-Cre;*Hes1*^*CKO/CKO*^ and Rax-Cre;*Hes*^*TKO*^ mutants with mispositioned Pax2 domains (arrows in Figs 4B,4C). Interestingly, Chx10-Cre;*Hes*^*TKO*^ eyes also have fewer Ccnd2+Pax2+ cells. Because *Hes1* encodes a transcriptional repressor, we presume the regulation of Ccnd2 is indirect. Once again, only Chx10-Cre;*Hes*^*TKO*^ retinal cells ectopically expressed Pax2 (Fig 4D), consistent with an expansion of ONH tissue in *Pax2*^*GFP*^ germline mutants [54]. Without *Pax2*, retinal cells are unable to lock-in a neural development program and express both RPC and ONH-specific markers [54]. This prompted us to ask whether *Hes*^*TKO*^ mutant RPCs, adjacent to the ONH, were similarly affected, using the ONH/OS marker *Vax1* [68-70](Fig 4E-H; n=3 replicates/ genotype). In Rax-Cre;*Hes1*^*CKO/CKO*^ and Rax-Cre;*Hes*^*TKO*^ eyes *Vax1* shifts down into the OS (arrows in Figs 4F, 4G). But only in Chx10-Cre;*Hes*^*TKO*^ eyes did the *Vax1* domain also extend in the opposite direction, into the retina (Fig 4H). These data suggest that sustained *Hes1* in the ONH helps lock-in the boundary with the retina, whereas multiple *Hes* genes in adjacent RPCs are necessary for maintaining neurogenic potential.

**Fig 4.**
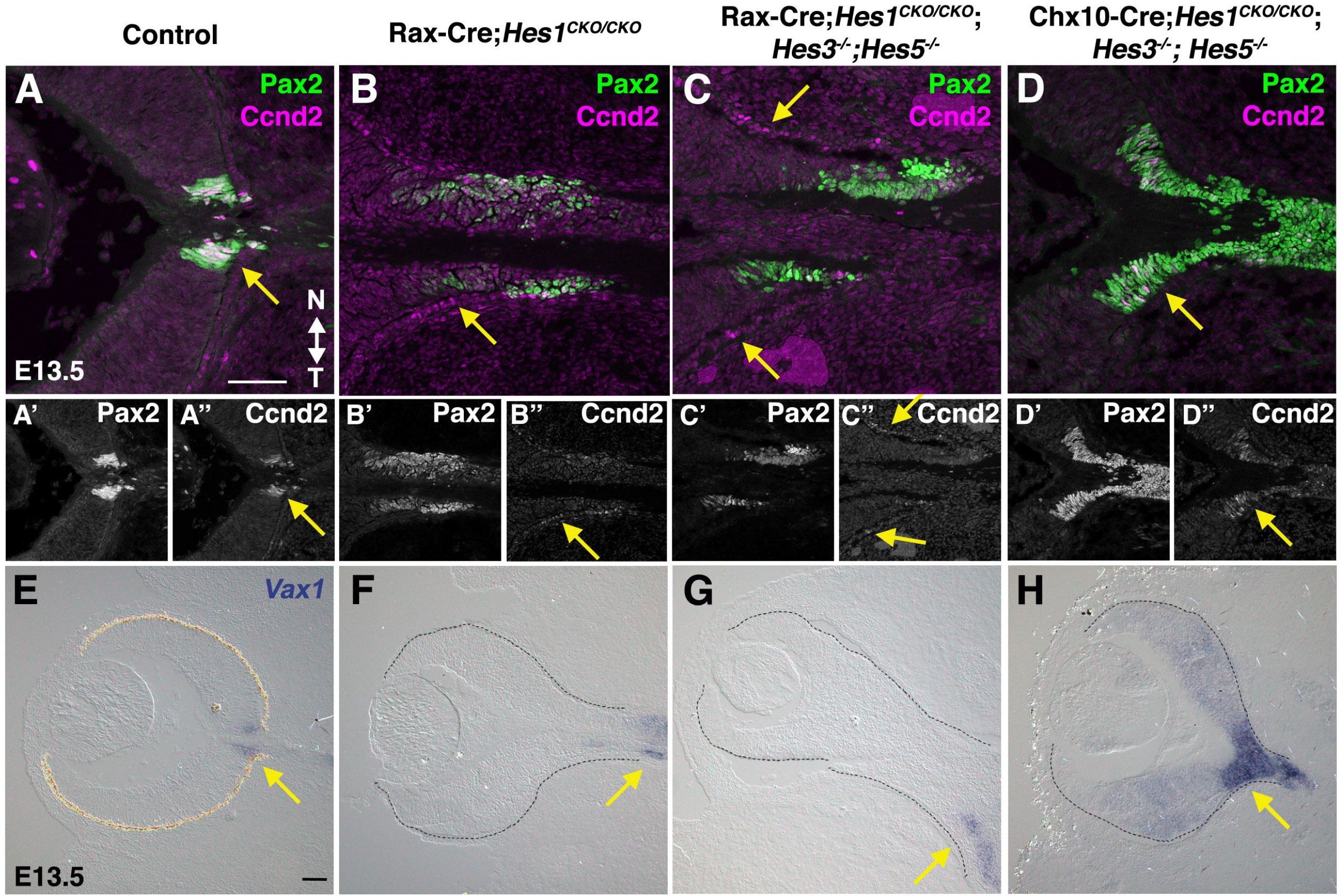
*Hes* gene mutant phenotypes at the retina-ONH boundary. A-D”) Pax2 + Ccnd2 immunolabeling at E13.5. Normally, Pax2 and Ccnd2 are coexpressed in ONH cells. In Rax-Cre;*Hes1*^*CKO/CKO*^ and Rax-Cre;*Hes*^*TKO*^ eyes, the Pax2 domain is elongated in the optic stalk, while Ccnd2 expression is dramatically downregulated in the optic stalk or mislocalized in the RPE (arrows in A,A”, B,B”, C,C”). Intriguingly, in Chx10-Cre;*Hes*^*TKO*^ eyes, both Pax2 and Ccnd2 domains expanded into the optic cup (arrow D”). (E-H) *Vax1* mRNA expression in the ONH/OS (arrows). Eyes in F-H are albino and the retina is outlined with dotted lines. The *Vax1* domain was shifted proximally in Rax-Cre;*Hes1*^*CKO/CKO*^ and Rax-Cre;*Hes*^*TKO*^ eyes, but is derepressed into the retina of Chx10-Cre;*Hes*^*TKO*^ eyes. N = 3 biologic replicates/genotype.

### Notch pathway regulation of RPC growth, death and onset of neurogenesis

Throughout the CNS, Notch signaling stimulates progenitor cell growth and blocks neurogenesis. We expected proliferation to be reduced in these six mutants and confirmed this by quantifying PhosphoHistone H3 (PH-H3) expression within G_2_ and M-phase cells (Figs 5A-5G, 5O). Both *Rbpj* mutants had the biggest reduction in mitotic cells. There was also a modest loss of PH-H3+ cells in *Hes*^*TKO*^ mutants for the Chx10-Cre driver, but not Rax-Cre. The opposite outcome was seen in *ROSA*^*dnMAML-GFP/+*^mutants. Reduced RPC proliferation is common to all mutants, although the age of phenotypic onset differs (Suppl Table 1).

**Fig 5.**
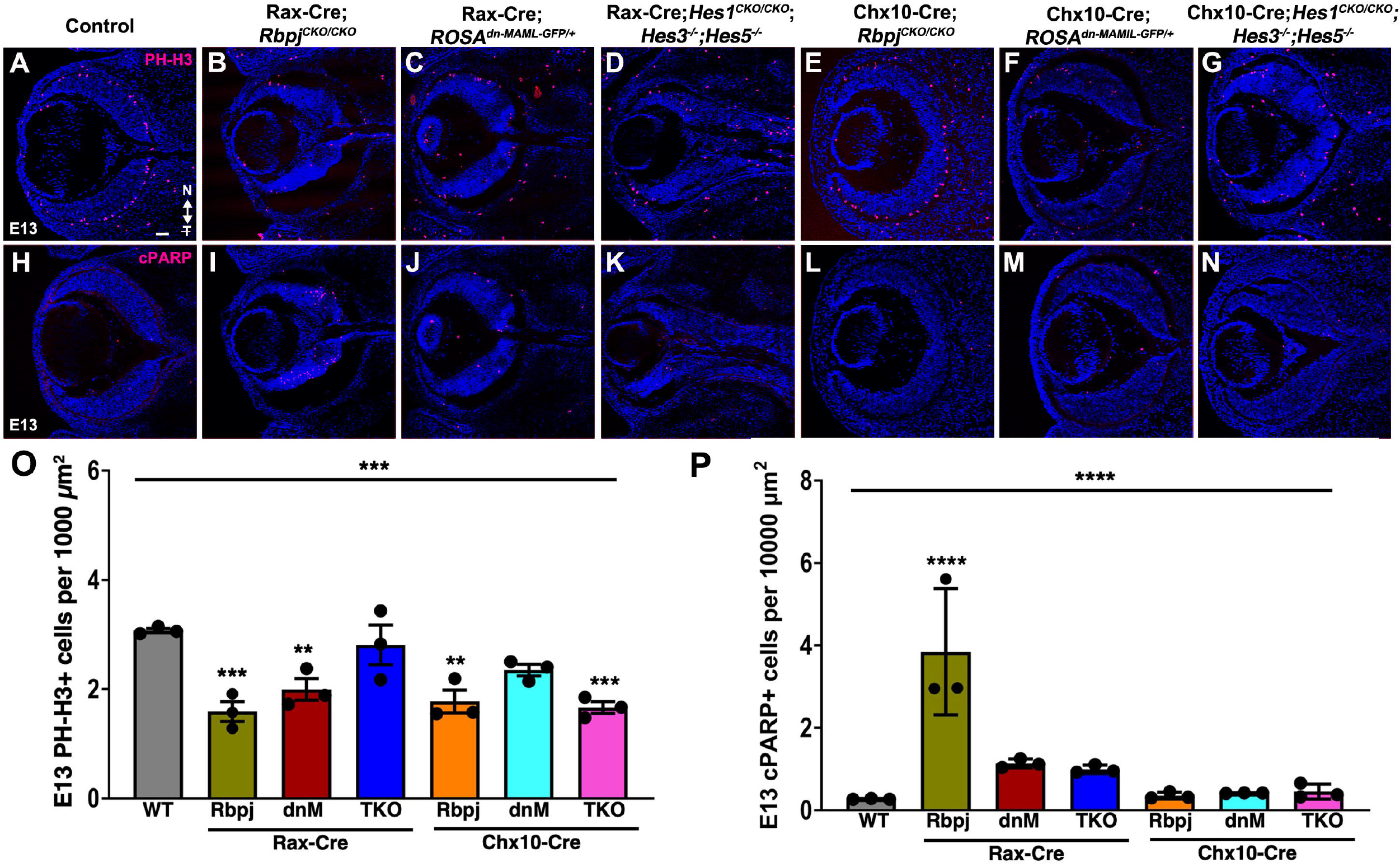
All E13.5 mutants have reduced proliferation, but only *Rbpj* mutants have excess apoptosis. (A-G) M-phase RPCs labeled with anti-PhosphoHistone-H3 (PH-H3) in red, DAPI in blue. (H-N) E13.5 cPARP+ apoptotic retinal cells in red, DAPI in blue. (O,P) Graphs display individual replicate data points, the mean and S.E.M; Significant Welch’s ANOVA, plus pairwise comparisons to wild type (****p< 0.0001, ***p<0.001, **p<0.01). All panels are oriented nasal up (noted in A); n = ≥2 sections from 3 biological replicates/genotype; scalebar in A = 50 microns.

In the E13-E16 retina, *Notch1, Rbpj* and *Hes1* mutants have a significant increase in apoptosis (Supplemental Table 1) [15, 17, 18, 38]. So, we used cPARP labeling to quantify dying cells among the six mutants (Figs 5H-5N, 5P). There was the anticipated increase in cPARP+ cells in E13.5 Rax-Cre;*Rbpj*^*CKO/CKO*^ mutants (Figs 5I, 5P), but all other genotypes were unaffected (Fig 5P). This suggests that Rax-Cre;*Hes*^*TKO*^ mutants rescued the apoptosis phenotype previously reported for *Hes1* single mutants [38]. This outcome could be attributed to either the loss of *Hes1* and *Hes5* coordinate regulation of target genes in RPCs, or inherent interactions between retinal and ONH tissues, which impacts cell viability.

The first postmitotic neurons appear in the central optic cup, and differentiation radiates outward from this point to the periphery. Loss of Notch signaling accelerates neurogenesis, with the first-born neurons quickly overproduced. Expression of the bHLH proneural factor *Atoh7* foreshadows the progression of neurogenesis from central to peripheral (Fig 6A)[71, 72]. Although *Atoh7* is not expressed by differentiated RGCs, its activity is required for their genesis [73-76]. Once differentiated, RGCs express neural-specific beta-tubulin (Tubb3) and the RNA-binding protein Rbpms, which is confined to RGCs within the retina [77, 78]. Both Rax-Cre;*Rbpj*^*CKO/CKO*^ and Rax-Cre;*Hes*^*TKO*^ eyes have dramatic overproduction of Rbpms+ and Tubb3+ RGCs at E13.5 and were missing Atoh7 in proximal retinal cells (Figs 6B, I, P; 6D, K, R). Ahead of the appearance of ectopic RGCs, Atoh7 was presumably precociously activated and downregulated, akin to what happens at E9.5 in *Hes1*^*-/-*^ mutants [47]. E13.5 Chx10-Cre;*Hes*^*TKO*^ eyes had a milder RGC phenotype (Figs 6G, 6N, 6U), but all other mutants were unaffected (Figs 6E, 6F, 6L, 6M, 6S, 6T). As is the case for Vsx2, we postulated proximal optic cup cells in Rax-Cre;*Rbpj*^*CKO/CKO*^ and Rax-Cre;*Hes*^*TKO*^ mutants were already differentiated. This is consistent with changes in Atoh7, Rbpms and Tubb3 expression (Figs 6B, 6D, 6I, 6K, 6P, 6R). Chx10-Cre;*Hes*^*TKO*^ eyes phenocopy this defect to a lesser extent (Figs 6N arrow, 6U). Interestingly both E13.5 Cre*;ROSA*^*dnMAML-GFP/+*^ eyes had clusters of mispositioned Rbpms+ RGCs in the temporal retina (Figs 6J, 6M arrow). At E16.5, all mutant retinas contained a vast excess of Rbpms+ or Tubb3+ RGCs (Figs 6V-6AA; 7U-7Z).

**Fig 6.**
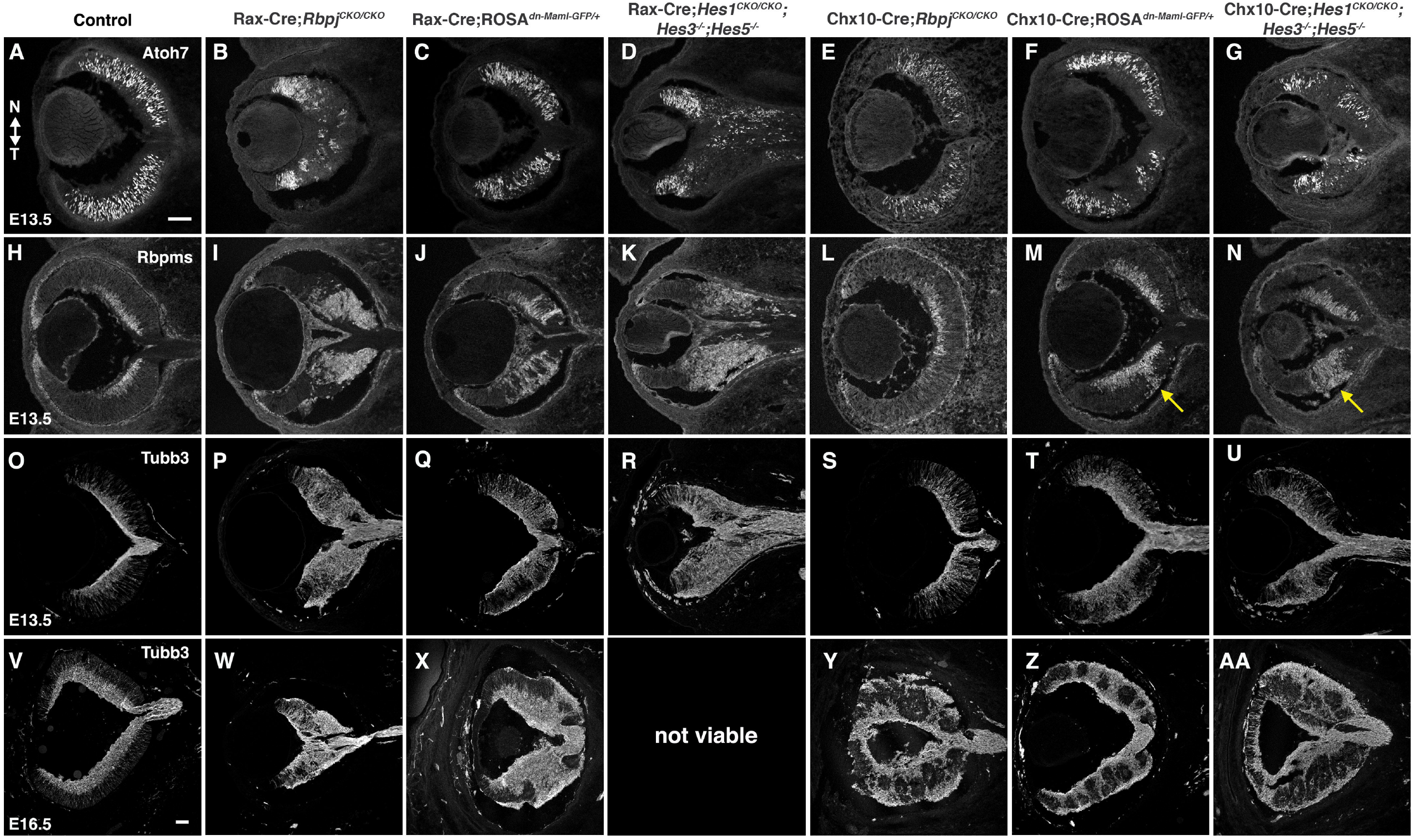
RGC neurogenic phenotypes among Notch pathway mutants. (A-G) Anti-Atoh7 labeling at E13.5 highlights reduction of retinal neurogenesis in Rax-Cre;*Rbpj*^*CKO/CKO*^ and Rax-Cre;*Hes*^*TKO*^ eyes, plus inappropriate Atoh7+ cells in the *Hes*^*TKO*^ optic stalk (D). (H-K) E13.5 Rbpms labeling shows clustered RGCs in the proximal optic cup for Rax-Cre;*Rbpj*^*CKO/CKO*^ and Rax-Cre;*Hes*^*TKO*^ eyes, and spreads down the optic stalk for the latter genotype (K). (L-N) Chx10-Cre;*ROSA*^*dnMAML*^ and Chx10-Cre;*Hes*^*TKO*^ eyes display excess, mispatterned RGCs (arrows) in the temporal retina. (O-U) E13.5 and (V-AA) E16.5 Tubb3+ retinal neurons are conspicuously increased in all Notch pathway mutants, regardless of Cre driver. All panels are oriented nasal up (noted in A); n = 3 biologic replicates/genotype scalebars in A,V = 50 microns.

### Notch pathway regulation of early photoreceptor cell fates

An archetypal retinal defect that arises after reduced Notch signaling is the appearance of retinal rosettes and an overproduction of Crx+ photoreceptors (Suppl Table 1) (Fig 7). *Hes1* germline and conditional mutants display retinal rosettes, but uniquely have a significant loss of cone photoreceptors that could not be attributed to developmental delay [15]. This incongruity raises questions about how the Notch pathway operates downstream of the ternary complex during photoreceptor genesis. In *Hes1* single mutants, *Hes5* mRNA is expanded (Fig 1G), implying that cone photoreceptor genesis might become blocked by ectopic *Hes5*. This is consistent with the exclusion of Hes5-GFP from Rxrg+ cone photoreceptor cells [49]. Therefore, we reasoned that simultaneous loss of multiple *Hes* repressor genes would rescue cone genesis, but might produce ectopic cones if fate choice were uncoupled from Notch pathway regulation of RPC proliferation versus differentiation.

**Fig 7.**
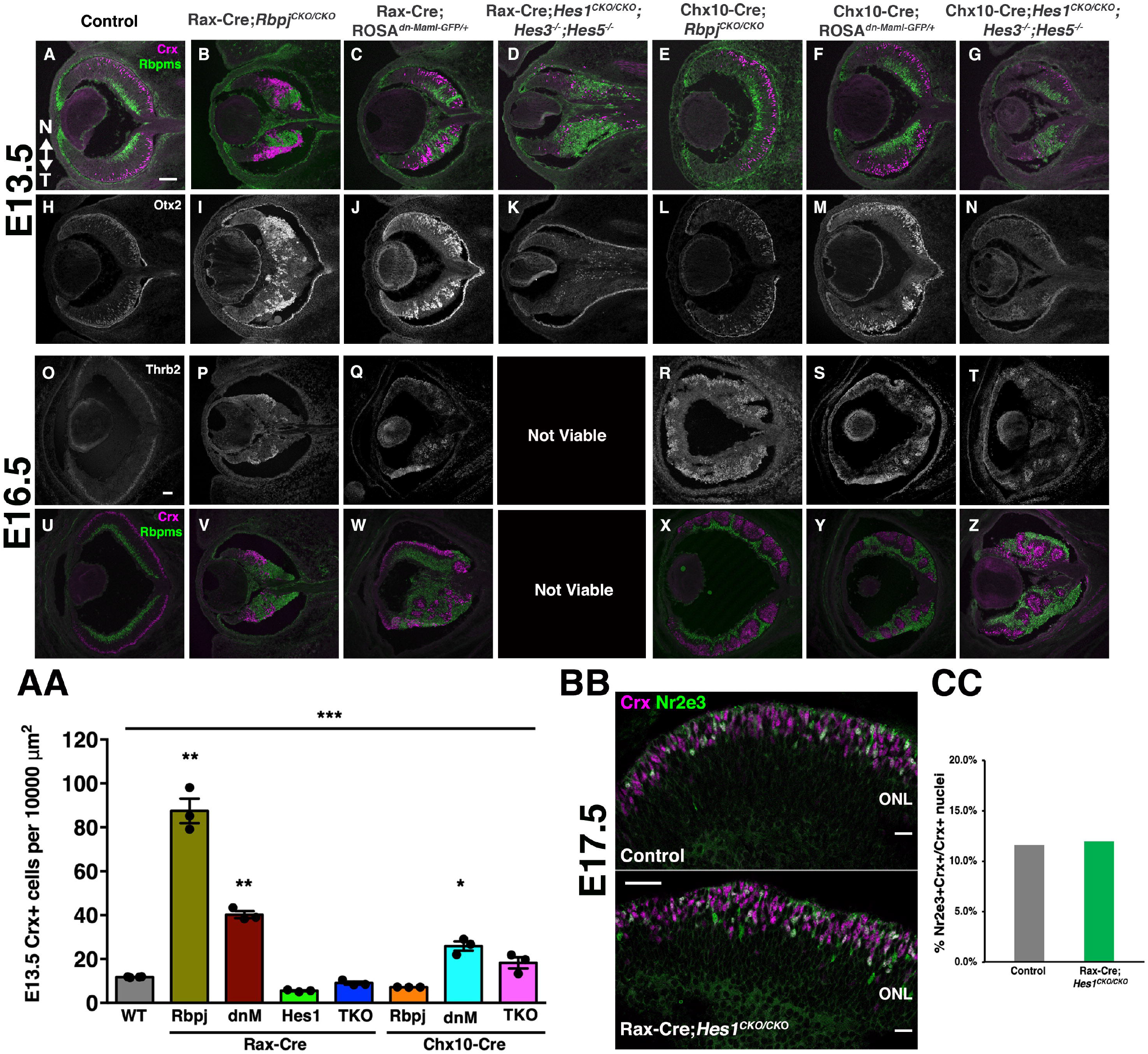
Notch pathway genes do not regulate embryonic photoreceptor fates equally. (A-G) Crx-Rbpms double labeling at E13.5 reveals early mispatterning of mutant retinas with varying shifts in the numbers and distribution of Rbpms+ RGCs and Crx+ photoreceptors. (H-N) Otx2+ cells at E13.5. Otx2+ cells are elevated and abnormally clustered when Rax-Cre was used to delete *Rbpj* or to drive dnMAML expression. However, there are conspicuously fewer Otx2+ cells in Rax-Cre;*Hes*^*TKO*^ eyes. These defects were much milder for the Chx10-Cre induced alleles (E-G, L-N). (O-T) E16 immunostaining for Thrb2. There are fewer Thrb2+ cones in all genotypes, except Chx10-Cre;*Hes*^*TKO*^. (U-Z) Crx-Rbpms colabeling suggests most retinal cells are RGCs or photoreceptors. Retinal rosettes are filled with Crx+ cells. (AA) Quantification of E13.5 Crx+ cells. Rax-Cre;*Rbpj* mutants have a unique increase in Crx+ cells. Graph displays individual replicate data points, the mean and S.E.M; Significant Welch’s ANOVA, plus pairwise comparisons to wild type (**p<0.01, * p< 0.05). (BB, CC) No difference in the proportion of Nr2e3+ rods in the Crx lineage are seen between genotypes. Panels A=Z oriented nasal up, n = ≥2 sections from ≥3 biological replicates/genotype. Panels in (BB) oriented scleral up. (CC) Quantification from n = 11 whole retinal tile scanned composite image sections from 2 biologic replicates/genotype. Graph displays the mean and standard deviation. Scalebar in A, O = 50 microns, in BB = 20 microns.

To test these ideas, we directly compared the expression of early markers for the photoreceptor lineage in the Notch pathway allelic series. A large subset of embryonic RPCs expresses the transcription factor *Otx2* and are initially capable of producing five fates: cone, rod, amacrine, horizontal or bipolar neurons [79-81]. However, Otx2 is shut off relatively quickly in those cells that will adopt amacrine and horizontal fates. The remaining Otx2-lineage cells, which produce cones, rods and bipolar neurons [8], then activate the transcription factor Crx [82-85]. In *Notch1* and *Rbpj* mutants it was already known there are excess Otx2- and Crx-expressing cells [9-18](Suppl. Table 1), so we labeled our mutants with antibodies for Otx2 and Crx at E13.5 or E16.5 (Fig 7 and data not shown). First we re-confirmed that Rax-Cre;*Rbpj*^*CKO/CKO*^ mutants display ectopic Otx2+ and Crx+ cells (Figs 7B, 7I, 7AA), but noted that Chx10-Cre; *Rbpj*^*CKO/CKO*^ eyes were not different from controls (Figs 7E, 7AA). We attributed the latter outcome to Chx10-Cre mosaic activity and wild-type cell phenotypic rescue (Suppl Fig 2). We also verified that ectopic Crx+ cells in *Rbpj* mutants are Thrb2+cones (Figs 7O-T) and not precocious Nr2e3+ rods (not shown). We also found that both *ROSA*^*dnMAML-GFP/+*^ mutants weakly phenocopied Rax-Cre;*Rbpj*^*CKO/CKO*^ (Figs 7C, 7F, 7AA), but curiously, these photoreceptor rosettes were confined to the temporal side of E16.5 Rax-Cre;*ROSA*^*dnMAML-GFP/+*^ retinas (Fig 7W). Finally, E13.5 *Hes*^*TKO*^ mutant retinas contain normal proportions of Crx+ cells, regardless of Cre driver used (Figs 7D,7G, 7AA). We conclude that the loss of multiple *Hes* genes rescues the *Hes1* single mutant cone phenotype, but fails to phenocopy the rest of the Notch pathway (ectopic cones).

Previously E17.5 *Hes1*^*-/-*^ ex vivo retinal cultures were reported to contain rosettes, premature rod photoreceptor formation and reduced bipolar neurons [30]. We wished to revisit these outcomes, since we found that P21 *Hes1* conditionally mutant eyes have excess bipolar neurons [38]. Here the proportion of rods within the Crx+ cohort of E17 littermate control and Rax-Cre;*Hes1*^*CKO/CKO*^ retinal sections were quantified, by colabeling for Crx and Nr2e3, a transcription factor specifically present in nascent rods [86]. Nr2e3+ nuclei were present in the forming outer nuclear layer (ONL) for both genotypes (Fig 7BB), and retinal rosettes (not shown). However, the percentages of Nr2e3+Crx+ cells were identical between genotypes (Fig 7CC; n= 2 biologic replicates/genotype; per total Crx+ cells control = 10,661; RxCre;Hes1^CKO/CKO^ = 9,146 cells). Therefore, the loss of cones in *Hes1* mutants could not be attributed to accelerated rod genesis. We conclude that a proportion of the Otx2 lineage undergoes delayed terminal differentiation in the absence of *Hes1*, potentially adopting bipolar fates. Other possibilities include a greater acceleration of RGC development in *Hes1* mutants, and/or ectopic upregulation of *Hes5*, which does not happen in *Rbpj* or *Hes*^*TKO*^ mutant eyes.

When *Otx2* activity is blocked or removed, mutant cells switch from photoreceptor/bipolar to adopt amacrine/horizontal fates [79-81]. Thus, we reasoned Rax-Cre;*Rbpj*^*CKO/CKO*^ and Rax-Cre;*Hes1*^*CKO/CKO*^ single mutants can oppositely regulate RPC progression towards photoreceptor fates. Indeed, E13.5 *Rbpj* mutants have excess Otx2- and Crx-expressing cells [15, 18], whereas *Hes1* mutants have significantly fewer Otx2+ cells [Figure 6 in 38], yet an essentially normal cohort of Crx-expressing cells (Fig 7AA). To confirm this difference, we also evaluated Prdm1/Blimp1 expression, since it also acts downstream of Otx2 [87]. At, E13.5 Prdm1+ cells were quantified among all Rax-Cre induced mutants, plus Rax-Cre;*Hes1*^*CKO/CKO*^ single and Rax-Cre;*Hes1*^*CKO/CKO*^;*Hes3*^*+/-*^*;Hes5*^*+/-*^ mutants for better evaluation of the relative contributions of each *Hes* gene (Figs 8A-F, 8M). We found that only *Rbpj* mutants have excess Prdm1+ cells (Fig 8M). All other genotypes showed little to no reduction, most notably Rax-Cre;*Hes1*^*CKO/CKO*^ single mutants. Together these data suggest that the different activities of *Rbpj* and *Hes1* occur upstream of Otx2, but use distinct molecular mechanisms since opposite regulation of Crx and Prdm1 was revealed in each mutant.

**Fig 8.**
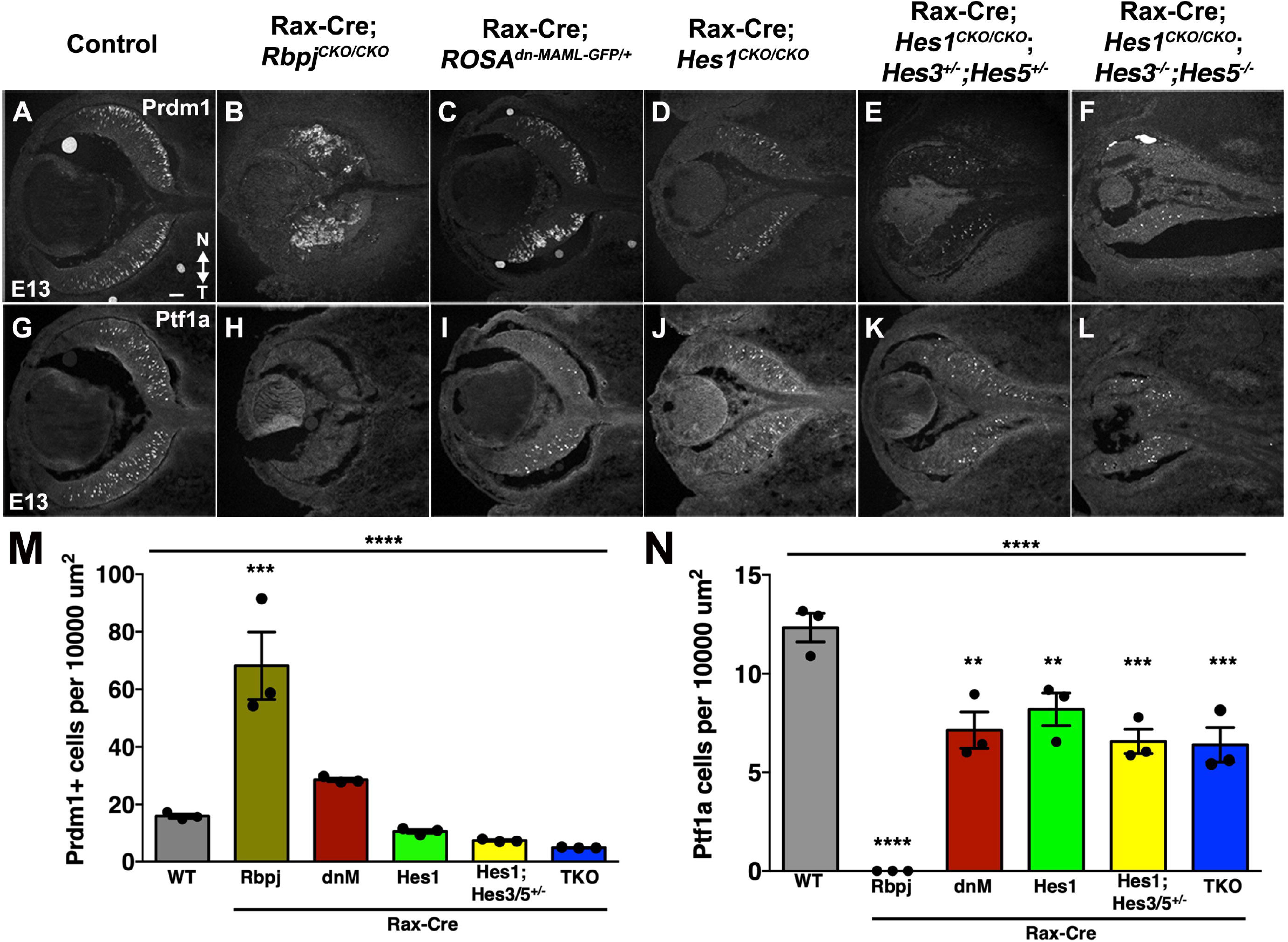
*Rbpj* and *Hes1* have opposing activities for photoreceptor versus amacrine fates. Prdm1/Blimp1 (A-F) and Ptf1a (G-L) labeling of E13.5 Rax-Cre-meditated deletion of Notch pathway genes. (M-N). Strikingly, only *Rbpj* mutants have excess Prdm1+ cells and a total loss of Ptf1a-expressing cells, whereas the loss of *Hes1* or multiple *Hes* genes partially rescues both phenotypes. For Rax-Cre;ROSA^dnMaml1-GFP/+^ retinas (where Hes expression is partially knocked down, see Fig 2), we note the partial rescue of Prdm1 and Ptf1a. (M,N) Graphs display individual replicate data points, the mean and S.E.M; Significant Welch’s ANOVA (****p< 0.0001; plus several pairwise comparisons to wild type (***p<0.001, ** p< 0.01). All panels oriented nasal up (noted in A; n = ≥2 sections from 3 biological replicates/genotype, scalebar in A = 50 microns.

Within the early Otx2 lineage, cells normally transiting to amacrine or horizontal fates downregulate Otx2 as they activate the transcription factor Ptf1a [reviewed in 88]. Subsequently, newly born amacrines activate Tfap2a/Ap2*α*. Ptf1a is both necessary and sufficient for amacrine and horizontal fates, and retinal cells that lose Ptf1a erroneously differentiate into RGCs and photoreceptors [89-91]. Without *Rbpj* we saw no Ptf1a+ cells or Tfap2a+ amacrines (Fig 8H,N), yet each mutant lacking *Hes1* had at best, a small reduction of Ptf1a, that probably reflects the diminished pool of RPCs (Fig 8N). The consequences of removing *Rbpj* on the amacrine pathway agree with previous studies (Suppl Table 1)[18, 89, 91], and further reinforce that Ptf1a expression depends on *Rbpj*.

## Discussion

The molecular mechanisms integrating Notch with other signaling pathways remain poorly understood. Here we directly compared the genetic requirements for ternary complex components and multiple *Hes* genes at the onset of retinal neurogenesis. We found that every gene acts to control the rate and timing of RPC proliferation. Interestingly, we also delineated pathway branchpoints, especially at the onset of photoreceptor neurogenesis.

### Hes genes in the developing eye

*Hes* genes are negative regulators of neuronal differentiation. They are critical for maintaining the proper number, age of appearance and spatial arrangement of each cell type. In the developing mammalian eye, *Hes1* and *Hes5* have been examined multiple times, using a variety of genetic tools [12, 15, 19, 30, 38, 40, 47, 49]. *Hes1* has multiple activities. It maintains optic vesicle and cup growth when expressed at high levels, it sets the tempo of neural differentiation throughout retinogenesis while in oscillatory expression mode, and in ONH and OS cells it acts via sustained expression to promote astrocyte development. By contrast, *Hes5* may only act on its own during postnatal Müller gliogenesis, although *Hes1* activity is required here too [19, 92]. In other areas of the CNS, *Hes3* is active during oligodendrocyte maturation and interacts with STAT3-Ser and Wnt signaling pathways prior to the initiation of myelination [93, 94]. Given that *Hes3* mRNA is undetectable in the embryonic retina, we propose it is relatively more important postnatally, possibly for retinal astrocyte migration, and/or astrocyte myelination of the optic nerve.

### Making and keeping the retinal-glial boundary

The retinal ONH/OS possesses many of the characteristics of the brain isthmus, which is comprised of slowly proliferating cells that undergo little to no neurogenesis and act as a signaling hub for adjacent neural tissues [reviewed in 27, 95]. Consistent with this idea, we found *Hes1* is required for Ccnd2 expression, which is associated with prolonged cell cycles. Both the ONH and isthmus require the transcription factors *Hes1* and *Pax2*. In the eye, loss of either gene allows the retina to encroach and displace the ONH. This expansion might be due to a failure to effectively shift from fast to slow cycling kinetics or from ectopic *Hes5* expression. In this developmental context, the loss of multiple *Hes* genes was essentially the same as that of *Hes1* alone. Thus, sustained *Hes1* is likely sufficient for ONH formation and maintenance. Our data do not support a role for Notch signaling in the ONH/OS, since both *Rbpj* and *ROSA*^*dnMAML-GFP/+*^ retain an attenuated, but recognizable ONH and ONH cells retain sustained Hes1 expression (Fig 9A).

**Fig 9.**
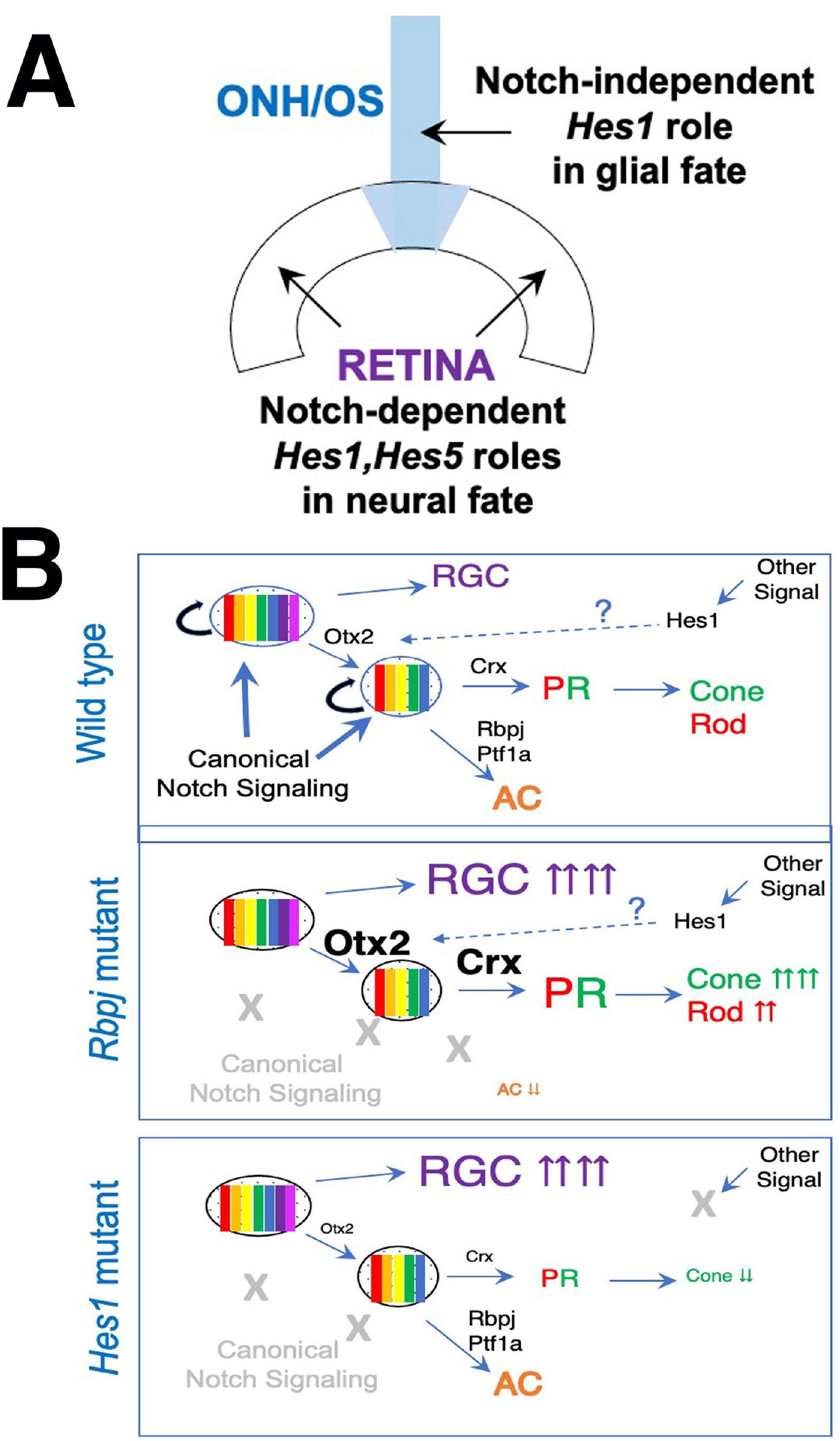
Roles for Notch signaling and potential integration points with other genetic pathways in the prenatal retina. (A) Maintenance of retinal versus ONH/OS territories: Sustained Hes1 expression in the ONH/OS does not require ternary complex gene activities (Notch-independent), while oscillating *Hes1* (and potentially *Hes5*) within RPCs maintain neuronal potential. (B) Contribute to cell type diversity: Notch signaling regulates neuron versus progenitor, whereas another signaling (X) provides specific feedback, through Hes1, about the onset of photoreceptor genesis.

In the ONH, sustained Hes1 expression must be regulated by other genetic pathways. A strong candidate is Shh pathway regulation of Hes1. Shh signaling performs an important feedback mechanism for RGCs to control their population size. Nascent RGCs secrete Shh, which instructs RPCs to remain mitotically active via direct binding of Gli2 to activate *Hes1* transcription [39, 96]. Moreover, at the optic vesicle stage of development, Shh diffuses from the ventral midline of the diencephalon to stimulate outgrowth of the optic cup and stalk [reviewed in 97]. Given that Wnt, Bmp and Retinoic Acid signals also regulate proximoventral optic cup and stalk outgrowth and specification [reviewed in 98], it is tantalizing to speculate that they do so by converging on Hes1 expression and/or activity that restricts the ONH to future glial lineages. However, it remains unresolved if the ONH is a signaling hub for the adjacent retina.

To delineate ONH versus retinal phenotypes, we used both Rax-Cre and Chx10-Cre drivers to test for functional redundancy of *Hes1* and *Hes5* during retinal neurogenesis. Unfortunately, the Chx10-Cre line could only produce a few robust outcomes, due to mosaic expression and the wild-type, nonautonomous rescue of some phenotypes. This problem was unanticipated since this driver was successfully used in previous genetic analyses of *Hes1, Rbpj* or *Neurog2* in the retina [15, 38, 99].

Because fewer and fewer Cre-GFP+ cells show activity over developmental time, we favor the ideas that this BAC transgene undergoes epigenetic silencing and/or wild type RPCs can outcompete the mutant ones over time. Nonetheless, we found that at E13.5 *Hes*^*TKO*^ eyes had abnormal ONH size and shape, and like *Pax2* mutants, ONH markers were expanded into the neural retina. We conclude that RPCs depend on the interplay of *Hes1* and *Hes5* to lock-in their neural fates. These studies also highlight the unequal retinal defects of *Rbpj* conditional mutants and dnMAML, a dominant-negative *Maml* allele, which disrupts ternary complex activity in other contexts; and the variable penetrance and severity of Rax-Cre versus Chx10-Cre drivers that are informative for future retinal studies.

### Notch-independent modes for regulating retinal histogenesis

An important goal of this study was to understand how precisely *Hes1* and *Hes5* activities mirror the Notch ternary complex, which can directly activate *Hes* gene transcription [reviewed in 4]. Because there are multiple ligands and Notch receptors expressed in the developing retina, we focused on the requirements for *Rbpj* (Fig 9B) and to a lesser extent *Maml*. There are three *Mastermind-like* paralogues (*Maml* genes), but germline mutant analyses failed to uncover individual gene functions during embryogenesis [reviewed in 100]. Subsequently, a dominant negatively acting isoform of MAML1 (dnMAML) was created, in which the MAML1 N-terminus forms ternary complexes with NICD and Rbpj, but cannot further interact with obligate transcriptional coactivators (e.g. p300, histone acetyltransferases) [42-45]. This has been a powerful tool in cancer biology and immunology research [42], but for retinal neurogenesis, we found that dnMAML is less effective at blocking Notch signaling. This might be attributed to differences in expression level (in vivo Cre-mediated induction here versus plasmid or viral delivery). However, we propose that Rax-Cre;*ROSA*^*dnMAML-GFP/+*^ mutants interfere with another gene pathway, due to a temporal-specific downregulation of Hes1 and *Hes5*, spatial restriction of photoreceptor rosettes, and Foxg1 mislocalization (Figs 2, 7, Suppl 3). In vitro proteomic studies support this idea, since dnMAML can bind to Gli and Tcf/Lef transcription factor proteins [101, 102]. This implies that *ROSA*^*dnMAML-GFP/+*^ retinal phenotypes may represent composite outcomes of simultaneously interfering with Notch, Shh and/or Wnt signaling.

*Rbpj* also has Notch-independent functions, the most common being its role in co-repressor protein complexes to silence transcription via DNA methylation [reviewed in 4]. Another activity is through Rbpj interactions with Ptf1a-E47 in a higher order PTF1 complex that has been studied in the pancreas, spinal cord and retina [88]. In the pancreas, PTF1 complexes can activate *Dll1*, suggesting this acts as a feedback loop from postmitotic to mitotic cells via Dll1 binding to Notch1 [103]. PTF1 directly antagonizes Notch signaling in a cell autonomous and dose-dependent manner, since Ptf1a and NICD bind to the same site on the Rbpj protein [88, 104]. Therefore, it is plausible that in the retina, PTF1 complexes divert RPCs from the photoreceptor lineage. We conclude that the *Rbpj* activity impacts early photoreceptor development in at least two ways. First, in the Notch ternary complex, it controls the timing of RPCs mitotic division versus differentiate, for example when RGCs or cone photoreceptors appear. Second, *Rbpj* prevents cells normally destined to become amacrines from erroneously developing as photoreceptors, via regulation of and independent physical interaction with Ptf1a.

These additional *Rbpj* and *Hes1* functions significantly complicate the interpretations of genetic datasets aimed at understanding how and where Notch signaling acts upstream of Otx2. For *Rbpj* mutants, the expansion of Otx2-, Crx-, and Prdm1-expressing cells and differentiated cone photoreceptors, at the expense of Ptf1a and amacrine neurons, fits current models of mutual exclusion for these factors [reviewed in 88]. However, the *Hes1* photoreceptor phenotype, fewer Otx2+ cells and cones, with much smaller impact on Crx-, Prdm1- or Ptf1a-expressing cohorts, further supports that *Hes1* regulation is a branchpoint for integrating information from other pathways. Alternatively, since *Hes1* mRNA and protein are dynamic, it will be critical to use live imaging and short-lived *Hes* reporters across these early stages of retinal development. Such dynamic expression is an inherent to the establishment of cellular heterogeneity and also can convey pulsatile feedback to other oscillating molecules like *Dll1, Neurog2* or *Ascl1* [28, 29], either upstream or downstream of Otx2, given there is a specific loss of Prdm1+ cells and rods in postnatal *Neurog2* mutants [99, 105]. Future studies that apply single cell imaging and sequencing modalities to remaining questions about when and where Notch signaling is required during retinal neurogenesis, particularly at the level of Otx2 regulation will be illuminating

## Materials and Methods

### Animals

Mouse strains used in this study are Hes5-GFP BAC transgenic line (*Tg(Hes5-EGFP)CV50Gsat/Mmmh* line; stock 000316-MU)[49, 106]; *Hes1*^*CKO*^ allele (*Hes1*^*tm1Kag*^) maintained on a CD-1 background [41]; *Rbpj* ^*CKO/CKO*^ (*Rbpj*^*tm1Hon*^) on a C57BL/6J background[9]; ROSA26^dnMAML-GFP^ (*Gt(ROSA)26Sor*^*tm1(MAML1)Wsp*^) maintained on a C57BL/6J background [42-45]; *Hes1*^*CKO/CKO*^;*Hes3*^*-/-*^;*Hes5*^*-/-*^ (*Hes1*^*tm1Ka*^)(*Hes3*^*tm1Kag*^) (*Hes5*^*tm1Fgu*^) triple homozygous stock, maintained on CD-1 and termed “ *TKO”* in this study [31, 41]; *Hes3*^*-/-*^;*Hes5*^*-/-*^ mice, derived from the triple stock; Chx10-Cre BAC transgenic line (*Tg Chx10-EGFP/cre;-ALPP)2Clc*; JAX stock number 005105) maintained on a CD-1 background [46]; and Rax-Cre BAC transgenic line (Tg(Rax-cre) NL44Gsat/Mmucd created by the GENSAT project [106], cryobanked at MMRRC UC Davis (Stock Number: 034748-UCD), and maintained on a CD-1 background. PCR genotyping was performed as described [1-9]. Conditional mutant breeding schemes mated one heterozygous Cre mouse to another mouse homozygous for GeneX conditional allele to create Cre;*GeneX*^*CKO/+*^ mice. The Cre;*GeneX*^*CKO/+*^ mice were used in timed matings with *GeneX*^*CKO/CKO*^ mice (see Suppl Table 2). The date of a vaginal plug was assigned the age of E0.5. All mice were housed and cared for in accordance with guidelines provided by the National Institutes of Health and the Association for Research in Vision and Ophthalmology, and were conducted with approval and oversight from the UC Davis Institutional Animal Care and Use Committee.

### Histology and Immunofluorescent Labeling

For histology, P21 eyes were dissected and fixed in 4% paraformaldehyde overnight at 4 °C and processed through standard dehydration steps and paraffin embedding. Four micron sections were stained with Hematoxylin and Eosin (H&E). For immunofluorescence, embryonic heads were fixed in 4% paraformaldehyde/PBS for 1 hour on ice, processed by stepwise sucrose/PBS incubations, and embedded in Tissue-Tek OCT. Ten micron frozen sections were labeled as in [107] with primary and secondary antibodies listed in Suppl Tables 3 and 4. Nuclei were counterstained with DAPI.

### RNA *in situ* hybridization

DIG-labeled antisense riboprobes were synthesized from mouse *Hes5* [49], and mouse *Vax1* [68] cDNA templates. In situ probe labeling, cryosection hybridizations and color development were performed using published protocols [71, 108].

### Microscopy and Statistical analysis

Histologic and in situ hybridization sections were imaged with a Zeiss Axio Imager M.2 microscope, color camera and Zen software (v2.6). Antibody-labeled cryosections were imaged using a Leica DM5500 microscope, equipped with a SPEII solid state laser scanning confocal and processed using Leica LASX (v.5) plus Navigator tiling subprogram, FIJI/Image J Software (NIH) and Adobe Photoshop (CS5) software programs. All images were equivalently adjusted for brightness, contrast, and pseudo-coloring. At least 3 biologic replicates per age and genotype were analyzed for each marker, and at least 2 sections per individual were quantified via cell counting or tissue area measurements. Sections were judged to be of equivalent depth by presence of or proximity to the optic nerve. For E17 retinal sections, 11 tile scanned retinal sections from 2 biologic replicates/genotype were quantified. Marker+ cells in tissue sections were counted using the count tool in Adobe Photoshop CS5 and statistical analyses performed using Prism (GraphPad v9) or Excel (v16.16.11) software, with p-values determined with one-way ANOVA and pair-wise Dunnett’s test. A p-value less than 0.05 was considered statistically significant.

## Acknowledgements

The authors thank Ryoichiro Kageyama, Tasuko Honjo, and Ivan Maillard for mutant mouse strains; Doug Forrest for Thrb2 antibody; Cheryl Craft for Opsin antibodies; Chris Wright for Ptf1a antibody; Brad Shibata and Paul FitzGerald for assistance with histology; Amy Riesenberg, April Bird, Kelly McCulloh for technical support. The authors thank Anna La Torre and Tom Glaser for thoughtful comments and members of the UCD Friday Eye Development group for feedback and discussion. This work was supported by the National Institutes of Health, National Eye Institute grants R01-EY024272 to JAB, R01-EY013612 to NLB and P30 EY012576 to UC Davis.

## Figure legends

**Suppl Table 1. Summary of Notch pathway mutant phenotypes in mouse retina**.

**Suppl Table 2. Recovery of mutant embryos/neonates at relevant stages of eye development**.

**Suppl Table 3. Validated primary antibody markers**.

**Suppl Table 4. Secondary antibody reagents**.

**Suppl Fig 1. *Hes3***^***-/-***^***;Hes5***^***-/-***^ **double mutants have no discernible eye phenotypes**. A,B) Number and pattern of Pou4f+ RGCs is unaltered. C,D) Pax6+ RPCs and mitotic Ccnd1+ cells are unaffected. E,F) Cdkn1b+ postmitotic RGCs and Sox9+ RPCs, RPE and ONH cells are the same between control and double mutants. G-J) Adult Müller glia, labeled with Sox9 (G,H) or Rlpb1/CRALBP (I,J) are also normal. All panels are vitreal down, scleral up; n = 4 biologic replicates/genotype; scalebar in A, E = 20 microns.

**Suppl Fig 2. Relative efficiencies of Rax-Cre versus Chx10-Cre BAC Tg drivers**. A-A’) Normal E13.5 expression patterns of Rbpj and Hes1. B-B”) Rax-Cre induces a complete loss of *Rbpj* in optic cup, ONH and RPE Cre lineage (red in B’). This eliminates Hes1 in the cup and RPE, but not in the attenuated ONH (B”). C-D’) Anti-Rbpj and GFP labeling highlights Chx10-Cre-GFP mosaicism, with scattered GFP-neg retinal cells (red only nuclei in C, pink only in D). Chx10-Cre expression does not spread into the ONH (D). E-F’) In Chx10-Cre;*Rbpj* mutant littermates, Rbpj+ cells are dramatically reduced, although the Hes1 retinal domain is less effected (F). G-H’) At E16, Cre-GFP, Rbpj and Hes1 are normally coexpressed. I-J’) Proportionally bigger Cre-GFP-neg regions of Chx10-Cre;*Rbpj* mutant retinas express Rbpj. In J, islands of GFP+ mutant cells are surrounded by Hes1-expressing cells, which either did not undergo Cre recombination or are wild type cells that eventually outcompete and subsequently outnumber the mutant cells. n=3 biologic replicates/genotype; scalebar in A, C = 50 microns.

**Suppl Fig 3. Nasal-temporal patterning in *Notch* pathway mutants**. (A-G) At E13.5 Foxg1, normally localized to the nasal retina, is properly restricted among nearly all mutants. In Rax-Cre;*Hes*^*TKO*^ eyes (D), the Foxg1 domain expanded into the optic stalk, consistent with other RPC markers. It remained biased to the nasal portion of the retina and optic stalk. (H-M) At E16.5, all mutants have nasally-restricted Foxg1 expression, except Rax-Cre;*ROSA*^*dnMAMl1-GFP/+*^ retinas that have some Foxg1+ nuclei present on the temporal side and within the adjacent subretinal space (arrow in J). All panels oriented nasal up (noted in A; n = 3 biologic replicates/genotype; scalebar in A, H =50 microns.

