## Supplemental Table 1 for "Not all Notch pathway mutations are equal in the embryonic mouse retina"

**Supplemental Table 1. Summary of mouse Notch signaling pathway retinal phenotypes**

| Mouse Gene | Mutation | Progenitors | Apoptosis | Retinal Ganglion Cells Neurons | Photoreceptors | Other Postnatal Cell types | Morphologic defects | Citation |
|---|---|---|---|---|---|---|---|---|
| **Ligands**[1] | | | | | | | | |
| *Dll1* | Conditional α Cre | ↓ | NT | ↑ **RGCs** and Neurons | **Cones** Normal **Rods** NT | Normal | R | Riesenberg et al 2016 |
| *Dll1* | Conditional Chx10 Cre | ↓ | ↑ | ↑ **RGCs** and Neurons | NT | NT | R OLM | Rocha et al 2009 |
| *Dll4* | Conditional Six3 Cre FoxN4 Cre | ↓ | ↑ E14 | **RGCs** NT ↑ Neurons | ↑ **Cones** E14.5 ↑ **Rods** >P0 | ↓ **AC** ↓ **BP** ↓ **MG** | Thinner retina R | Luo et al 2012 |
| *Jag2* | Germline Perinatal lethal | Normal | NT | Normal E14.5 | Not tested | Not tested | Normal E14.5 | Brown lab unpublished |
| **Receptors**[1] | | | | | | | | |
| *Notch1* | Germline Lethal ≥E10 | NA | NA | NA | NA | NA | NA | de la Pomp et al 1997 |
| *Notch1* | Conditional α Cre | ↓ | ↑ E14 | ↓ **RGCs** and Neurons | ↑ **Cones** E13.5 ↑ **Rods** ≥ P0 | NT | M, R | Yaron et al 2006 |
| *Notch1* | Conditional Chx10 Cre | ↓ | NT | ↓ **RGCs** and Neurons | ↑ **Cones** ≥E13.5 | NT | M, R | Jadhav et al 2006 |
| *Notch1* | Conditional ≥ P0 viral Cre | NA | NT | NA | ↑ **Rods** | ↓ **BP** ↓ **MG** | R | Jadhav et al 2006 Mizeracka et al 2013 |
| *Notch3* | Germline | Normal | NT | Normal | Normal | NT | NT | Maurer et al 2014 |
| *Notch1* *Notch3* | Conditional (α) Germline | ↓ | NT | ↑ **RGCs** and Neurons | ↑ **Cones** E13.5 | NT | NT | Maurer et al 2014 |

| | | | | | | | | |
|---|---|---|---|---|---|---|---|---|
| **Ternary complex** | | | | | | | | |
| *Rbpj* | Germline<br>Lethal ≥E8-9 | NA | NA | NA | NA | NA | NA | de la Pomp et al 1997 |
| *Rbpj* | Conditional<br>α Cre | ↓ | ↑ E16 | ↑ **RGCs** and Neurons | ↑ **Cones** E13.5<br>↑ **Rods** ≥ P0 | ↓ **BP**<br>↓ **MG** | M, R | Riesenberg et al 2009 |
| *Rbpj* | Conditional<br>Chx10 Cre | ↓ | ↑ E14 | ↑ **RGCs** and Neurons | ↑ **Cones** ≥ P0<br>↑ **Rods** ≥P0 | ↓ **AC** ↓ **BP**<br>↓ **MG** | M, R<br>Laminar defect | Zheng et al 2009 |
| *Maml1* | Germline | NT | NT | NT | NT | NT | Normal | Oyama 2007<br>Wu 2007 |
| *Maml3* | Germline | NT | NT | NT | NT | NT | Normal | Oyama et al 2011 |
| *Maml1*<br>*Maml3* | dbl Germline<br>Lethal ≥E10 | NA | NA | NA | NA | NA | NA | Oyama et al 2011 |
| **Effectors** | | | | | | | | |
| *Hes1* | Germline<br>Lethal ≥E15 | ↓ | NT | **RGCs** NT<br>↑ Neurons | **Cones** NT<br>Explants ↑ **Rods** | Explants<br>↓ **BP** ↓ **MG** | M, R<br>ONH | Tomita et al 1996 |
| *Hes1* | Germline<br>Lethal ≥E15 | ↓ | NT | ↑ **RGCs** and Neurons | NT | Explants<br>↓ **MG** | M, R<br>OLM; ONH | Takatsuka et al 2004 |
| *Hes1* | Germline<br>Lethal ≥E15 | ↓ | NT | ↑ **RGCs** and Neurons | NT | NT | M | Lee et al 2005 |
| *Hes1* | Conditional<br>Rax Cre<br>Chx10 Cre | ↓ | ↑ E16 | ↑ **RGCs** and Neurons | NT | ↑ **BP**<br>↓ **MG** | M, R<br>OLM<br>ONH | Bosze et al 2019 |
| *Hes3* | Germline | NT | NT | NT | NT | NT | Normal | Hirata et al 2001 |
| *Hes5* | Germline | Normal | Normal | NT | NT | ↓ **MG** | Normal | Hojo et al 2000 |

[1] *Jag1* and *Notch2* genes are required for lens fiber cell differentiation and ciliary body/iris formation; *Notch2* is required in the RPE

Abbreviations: NT = not tested; NA = not applicable; M = microphthalmia; R = retinal rosettes; RGC = retinal galnglion cells; AC = amacrines; BP = bipolars; MG = Muller glia; OLM = discontinuities in Outer Limiting Membrane; ONH = Expansion of retina into Optic Nerve Head/optic stalk
