## Supplemental Table 2 for "Not all Notch pathway mutations are equal in the embryonic mouse retina"

**Supplemental Table 2. Mendelian inheritance of evaluated Notch pathway mutant alleles.**

| Mouse mating | Expected Mutant Ratio | E10.5-E11 | E13.5 | E16.5 | P0 | P21 |
|---|---|---|---|---|---|---|
| Chx10Cre;*Rbpj*$^{CKO/+}$ **X** *Rbpj*$^{CKO/CKO}$ | 25% | 22%<br>14 / 65 from 6 litters | 36%<br>14 / 39 from 4 litters | 22%<br>11 / 51 from 5 litters | 23%<br>3 / 13 from 1 litter | 20%<br>3 / 15 from 6 litters |
| Rax-Cre;*Rbpj*$^{CKO/+}$ **X** *Rbpj*$^{CKO/CKO}$ | 25% | 24%<br>8 / 34 from 10 litters | 9%<br>3 / 33 from 6 litters | 31%<br>5 / 16 from 5 litters | 28%<br>1 / 6 from 2 litters | 30%<br>3 / 10 from 2 litters |
| Chx10Cre;*Hes* triple het [a] **X** *Hes* triple [b] | 12.5% [c] | No data | 21%<br>19 / 89 from 12 litters | 8%<br>4 / 52 from 11 litters | 14%<br>6 / 42 from 4 litters | 16%<br>14 / 96 from 6 litters |
| Rax Cre;*Hes* triple het [a] **X** *Hes* triple [b] | 12.5% [c] | 8%<br>6 / 75 from 14 litters | 14%<br>9 / 66 from 10 litters | Not viable<br>0 / 10 from 3 litters | No data | No data |
| Chx10Cre **X** *ROSA*$^{dnMaml1\text{-}GFP/\ dnMaml1\text{-}GFP}$ | 50% | 55%<br>2 / 36 from 5 litters | 54%<br>13 / 24 from 3 litters | 66%<br>10 /15 from 2 litters | 50%<br>3/6 from 1 litter | 50%<br>4/8 from 1 litter |
| Rax-Cre **X** *ROSA*$^{dnMaml1\text{-}GFP/\ dnMaml1\text{-}GFP}$ | 50% | 58%<br>21 / 36 from 6 litters | 52%<br>14 / 27 from 6 litters | 55%<br>11 / 20 from 3 litters | 44%<br>12 / 27 from 5 litters | 43%<br>3 / 7 from 1 litter |

[a] *Hes* triple het = *Hes1*$^{CKO/+}$;*Hes3*$^{KO/+}$;*Hes5*$^{KO/+}$

[b] *Hes* triple (Hes$^{TKO)}$ = *Hes1*$^{CKO/CKO}$;*Hes3*$^{KO/KO}$;*Hes5*$^{KOKO}$

[c] *Hes3* and *Hes5* genes <1Mb apart. Mutant alleles segregate together with no recombinants found by PCR genotyping (n = 52 litters)
