## Supplemental Table 3 for "Not all Notch pathway mutations are equal in the embryonic mouse retina"

**Supplemental Table 3: Validated Primary antibodies.**

Authentication methods used: 1: Antibody identification at http://antibodyregistry.org; 2: Genetic validation method; 3: Comparison with known published patterns in the literature; 4: Co-labeling of tagged protein with endogenous protein.

| Antigen | Source | Host | Catalog # (clone) | Dilution Used | Application | Authentication | Antibody ID | References (PMID) |
|---|---|---|---|---|---|---|---|---|
| Ap2alpha Tfap2a | DSHB | Mouse clone 3B5 | 3B5 | 1:50 conc sup | IF | 1,2,3 | AB_528084 | >10 |
| Atoh7 | Novus | Rabbit | NBP1-88639 Lot R12065 | 1:500 | IF | 1 2 3 4 | AB_11034390 | 29225067 |
| RFP-Biotin | Abcam | Rabbit | AB34771 | 1:1000 | IF | 1 3 | AB_777699 | 26587807 28641110 28817801 |
| Crx | Sigma/Prestige | Rabbit | HPA036762 Lot # R34426 | 1:1000 | IF | 1 2 3 4 | AB_10673663 | 31949107 |
| Cdkn1b* (p27) | BD/Invitrogen | Mouse | clone 57 | 1:100 | IF | 1 2 3 | AB_1075241 | 16690048 |
| Cyclin D1 (Ccnd1) | Neomarkers / Lab Vision | Rabbit | RM-9104-S1 Clone: SP4 | 1:100 | IF | 1 3 | AB_149913 | 15166673 |
| Cyclin D2 (Ccnd2) | Santa Cruz | Rat | SC-452 Clone 34B1-3 | 1:200 | IF | 1 2 3 | AB_627350 | 27957530 29533784 |
| Arr3 cone Arrestin | EMD Millipore | Rabbit | AB15282 | 1:700 | IF | 1 3 | AB_11210270 | 26453550 25798616 24651551 |
| cPARP (Mus-specific) | Cell Signaling | Rabbit | 9544 | 1:500 | IF | 1 3 | AB_2160724 | 27353360 30581080 31000436 |
| Foxg1 | Abcam | Rabbit | AB196868 | 1:1000 | IF | 1 3 | AB_2892604 | 32024767 |
| GFP | Aves | Chicken | GFP-1020 | 1:3000 | IF | 1 3 | AB100000240 | >100 in Registry |
| Glutamine Synthetase | EMD Millipore | Mouse | MAB302 (Clone GS-6) | 1:1000 | IF | 1 3 | AB_309678 | 26283925 26527153 25907681 |
| Hes1 | Cell Signaling | Rabbit | 11988 D6P2U | 1:500 | IF | 1 2 3 4 | AB_2728766 | 29681454 30059908 |
| Hes1 | Brown lab | Rabbit | N/A | 1:500 | IF | 2 3 4 | No data | 30059908 28675662 25100656 19828801 |
| Jagged1* (C20) | Santa Cruz | Goat | SC-6011 | 1:500 | IF | 1 2 3 4 | AB_649689 | 19389370 30890522 |
| Lhx2 | GENETEX | Rabbit | GTX129241 | 1:500 | IF | 1 3 | AB_2783558 | 31135891 |
| Mitf | ThermoFisher | Mouse IgG1 | MS-771-P1 | 1:100 | IF | 1 3 | AB_141542 | 31949107 33428890 |
| Opn1mw | Cheryl Craft | Rabbit | N/A | 1:1000 | IF | 1 3 | No data | 12853434 |
| Opn1sw | Cheryl Craft | Rabbit | N/A | 1:1000 | IF | 1 3 | No data | 12853434 |
| Otx2-Biotin | R&D Systems | Goat | BAF1979 | 1:1000 | IF | 1 3 | AB_2157171 | 28089909 30048641 |
| Pou4f/ Brn3 * | Santa Cruz | Goat | SC 6026 | 1:50 | IF | 1 3 | AB_673441 | 18626943 21246546 21452196 + 12 more |
| Pax2 | Biolegend Covance | Rabbit | 901001 PRB-276P | 1:1000 | IF | 1 2 3 | AB_2565001 AB_291611 | 2043750 2073760 2247382 3065975 |
| Pax6 | Biolegend Covance | Rabbit | 901301 PRB-278P | 1:1000 | IF | 1 2 3 4 | AB_2565003 AB_291612 | 28132835 + 24 more >40 in Registry |

| | | | | | | | | |
|---|---|---|---|---|---|---|---|---|
| Pax6 | Santa Cruz | Mouse ($IgG_{1}$) | SC-32766 (clone AD2.38) | 1:50 | IF | 1 2 3 | AB_628107 | 25100656 |
| Phospho Histone H3 | EMD Millipore | Rabbit | 06-570 Lot# DAM 1416518 | 1:500 | IF | 1 3 | AB_310177 | 17447250 18205207 18273885 >70 in registry |
| Prdm1/ Blimp1 | Santa Cruz | Rat | sc-47732 (clone 6D3) | 1:100 | IF | 1,2,3 | AB_628168 | 30184502 39463012 30850343 |
| Prox1 | Chemicon/Millipore | Rabbit | AB 5475 | 1:2000 | IF | 1,2,3 | AB_177485 | 17183554 +10 others |
| Ptf1a | Chris Wright | Rabbit | N/A | 1:2000 | IF | 1,2,3 | No data | 17075007 |
| Rax | TaKaRa | Rabbit | M228 | 1:1000 | IF | 3 | No data | 2911759 31949107 |
| Rbpj | Cosmo Bio Japan | Rat | SIM-2ZRBP1 Clone T6709 | 1:100 | IF | 1 2 3 | No data | 22275127 29977079 30059908 |
| Rbpms | PhosphoSolutions EMD Millipore | Guinea Pig | 1832-RPBMS ABN1376 | 1:500 | IF | 1 3 | AB_2492226 AB_2687403 | 25631988 27391320 +10 others |
| Rho | EMD Millipore | Mouse clone 1D4 | MAB5356 | 1:1000 | IF | 1 3 | AB_11215453 | 30401922 |
| Rlbp1 CRALBP | ThermoFisher | Mouse clone B2 | MA1-813 | 1:100 | IF | 1 3 | AB_2178528 | 11839540 11909957 +18 others |
| Rxrg* | Santa Cruz | Rabbit | SC-774 | 1:200 | IF | 1 3 | AB_2270041 | 39332629 30943581 |
| Six3 | Rockland | Rabbit | 600-401-A26 | 1:500 | IF | 1 2 3 4 | AB_11180063 | 30485816 |
| Sox2 | Millipore | Rabbit | AB5603 Lot # LV1395178 | 1:400 | IF | 1 2 3 | AB_2286686 | 16680766 >40 in registry |
| Sox9 | Millipore | Rabbit | AB5535 | 1:200 | IF | 1 3 | AB2239761 | 18626943 + 33 more |
| Thrb2 | Douglas Forrest NIH | Rabbit | N/A | 1:3000 | IF | 2 3 | No data | 19282790 |
| Tubb3 | Biolegend Covance | Rabbit | 802001 PRB-435P | 1:1000 | IF | 1 3 | AB_2564645 AB_291637 | 29247817 16874803 + others |
| Tubb3 | Biolegend Covance | Mouse ($IgG_{2A}$) | 801213 MMS-435P | 1:1000 | IF | 1 3 | AB_2564645 AB_291637 | 30695697 31209173 + others |
| Vsx2/Chx10 | Exalpha | Sheep | X1180P N-terminus | 1:500 | IF | 1 3 | AB_2314191 | 19827163 27565351 |
| Vsx2/Chx10 | Sigma/Prestige | Rabbit | HPA003436 | 1:500 | IF | 1 3 | AB_1078523 | none |
| ZO1 | Millipore | Rat | MABT11 Clone R40.76 | 1:500 | IF | 1 3 | AB_10616098 | 25078648, 23897660, 23991284 |

*: Discontinued; IF = Immunofluorescence; N/A = not applicable
