## Supplemental Table 4 for "Not all Notch pathway mutations are equal in the embryonic mouse retina"

**Supplemental Table 4. Secondary or Tertiary antibodies used in this study.**

| Antibody | Source | Host | Catalog Number | Dilution Used |
|---|---|---|---|---|
| Anti Chicken IgY Alexa 488 | ThermoFisher | Goat | A-11039 | 1:1000 |
| Anti-Sheep IgG Alexa 594 | Jackson Immuno. | Donkey | 713-585-147 | 1:500 |
| Anti-Guinea Pig IgG (H+L) Alexa 647 | ThermoFisher | Goat | A-21450 | 1:200 or 1:500 |
| Anti-Goat IgG Alexa 594 | ThermoFisher | Donkey | A11058 | 1:500 |
| Anti-Goat IgG (H+L) Alexa 647 | Jackson Immuno. | Donkey | 705-605-147 | 1:500 |
| Anti-Rabbit IgG Alexa 594 | ThermoFisher | Goat | A11037 | 1:500 |
| Anti-Rabbit IgG (H+L) Cy3 | Jackson Immuno. | Goat | 111-165-144 | 1:500 |
| Anti-Rabbit IgG Biotin-SP | Jackson Immuno. | Donkey | 711-005-152 | 1:1000 |
| Anti-Mouse IgG (H+L) Alexa 647 | ThermoFisher | Goat | A21236 | 1:200 |
| Anti-Mouse $IgG_1$ Alexa 647 | ThermoFisher | Goat | A212540 | 1:200 |
| Anti-Mouse IgG2a Alexa 647 | Jackson Immuno | Goat | 115-605-206 | 1:500 or 1:1000 |
| Anti-Rat IgG Alexa 594 | Jackson Immuno. | Donkey | 712-586-153 | 1:500 |
| Anti-Rat IgG (H+L) Alexa 647 | Jackson Immuno. | Donkey | 712-605-153 | 1:200 |
| Streptavidin Alexa 594 | Jackson Immuno. | Not applicable | S32356 | 1:1000 |
| Streptadivin Alexa 647 | Molecular Probes | Not aplicable | S32357 | 1:500 |
| DAPI 1 mg/ml | Sigma | Not applicable | D9542 | 1:500 |
