## Supplemental Figure 1 for "Not all Notch pathway mutations are equal in the embryonic mouse retina"

### Supplemental Figures

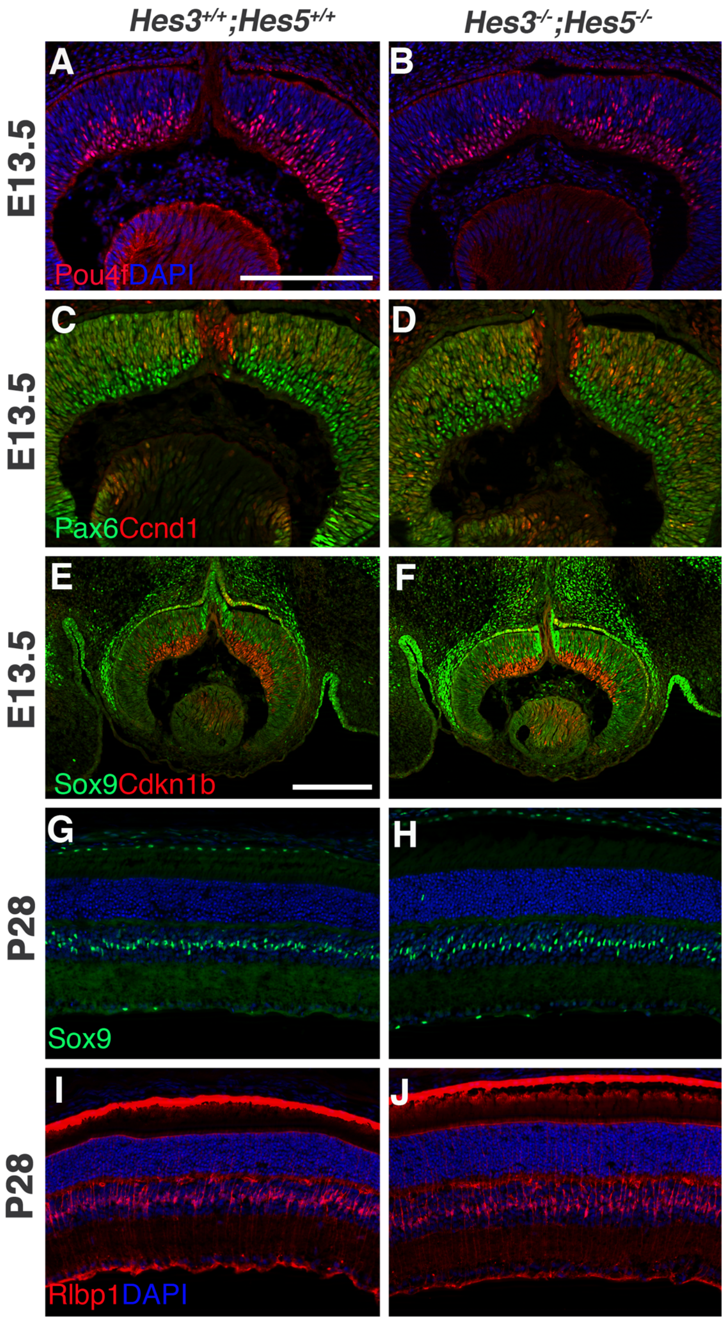


**Supplemental Figure 1.** *Hes3-/-;Hes5-/-* double mutants have no discernible eye phenotypes. A,B) Number and pattern of Pou4f+ RGCs is unaltered. C,D) Pax6+ RPCs and mitotic Ccnd1+ cells are unaffected. E,F) Cdkn1b+ postmitotic RGCs and Sox9+ RPCs, RPE and ONH cells are the same between control and double mutants. G-J) Adult Müller glia, labeled with Sox9 (G,H) or Rlpb1/CRALBP (I,J) are also normal. All panels are vitreal down, scleral up; n = 4 biologic replicates/genotype; scalebar in A, E = 20 microns.
