## Supplemental Figure 2 for "Not all Notch pathway mutations are equal in the embryonic mouse retina"

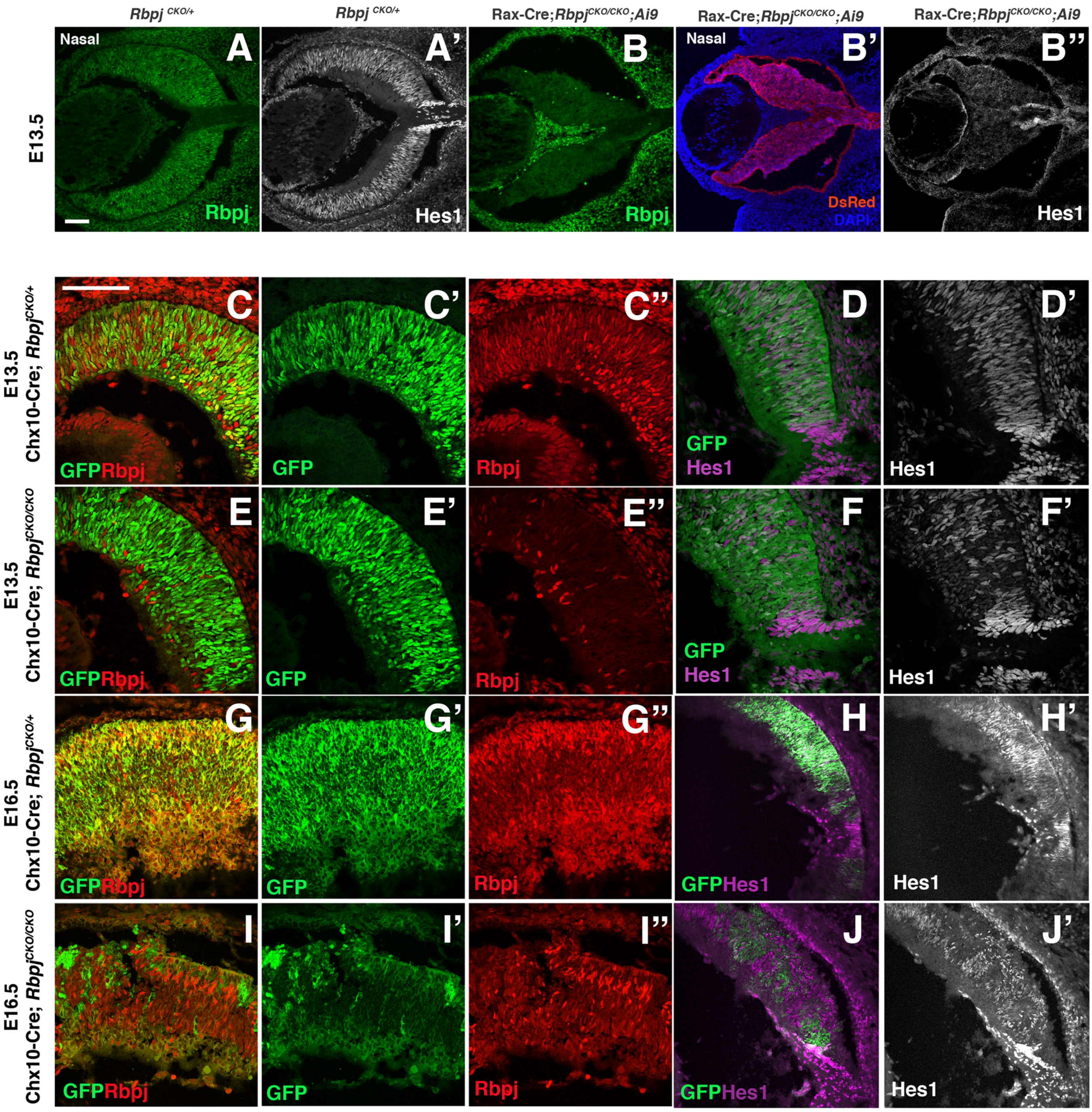


**Supplemental Figure 2**. Relative efficiencies of Rax-Cre versus Chx10-Cre BAC Tg drivers.
A-A') Normal E13.5 expression patterns of Rbpj and Hes1. B-B") Rax-Cre induces a complete loss of *Rbpj* in optic cup, ONH and RPE Cre lineage (red in B'). This eliminates Hes1 in the cup and RPE, but not in the attenuated ONH (B"). C-D') Anti-Rbpj and GFP labeling highlights Chx10-Cre-GFP mosaicism, with scattered GFP-neg retinal cells (red only nuclei in C, pink only in D). Chx10-Cre expression does not spread into the ONH (D). E-F') In Chx10-Cre;*Rbpj* mutant littermates, Rbpj+ cells are dramatically reduced, although the Hes1 retinal domain is less effected (F). G-H') At E16, Cre-GFP, Rbpj and Hes1 are normally coexpressed. I-J') Proportionally bigger Cre-GFP-neg regions of Chx10-Cre;*Rbpj* mutant retinas express Rbpj. In J, islands of GFP+ mutant cells are surrounded by Hes1-expressing cells, which either did not undergo Cre recombination or are wild type cells that eventually outcompete and subsequently outnumber the mutant cells. n=3 biologic replicates/genotype; scalebar in A, C = 50 microns.
