## Supplemental Figure 3 for "Not all Notch pathway mutations are equal in the embryonic mouse retina"

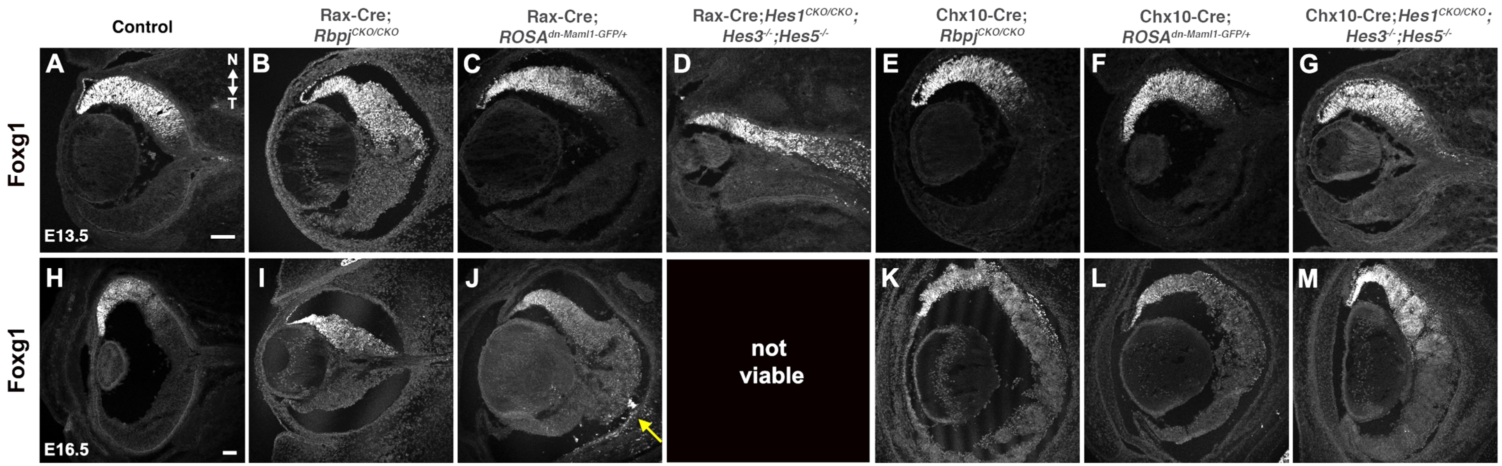


**Supplemental Figure 3.** Nasal-temporal patterning in Notch pathway mutants. (A-G) At E13.5 Foxg1, normally localized to the nasal retina, is properly restricted among nearly all mutants. In Rax-Cre;*Hes*$^{TKO}$ eyes (D), the Foxg1 domain expanded into the optic stalk, consistent with other RPC markers. It remained biased to the nasal portion of the retina and optic stalk. (H-M) At E16.5, all mutants have nasally-restricted Foxg1 expression, except Rax-Cre;*ROSA*$^{dnMAMl1\text{-}GFP/+}$ retinas that have some Foxg1+ nuclei present on the temporal side and within the adjacent subretinal space (arrow in J). All panels oriented nasal up (noted in A; n = 3 biologic replicates/genotype; scalebar in A, H =50 microns.
